# A large panel of chicken cells are invaded *in vivo* by *Salmonella* Typhimurium even when depleted of all known invasion factors

**DOI:** 10.1101/2020.11.17.386375

**Authors:** S. M. Roche, S. Holbert, Y. Le Vern, C. Rossignol, A. Rossignol, P. Velge, I. Virlogeux-Payant

## Abstract

Poultry are the main source of human infection by *Salmonella*. As infected poultry are asymptomatic, the identification of infected poultry farms is difficult. Controlling animal infections is thus of primary importance. As cell tropism is known to govern disease, our aim was therefore to identify the infected host-cell types in chicks and the role of the three known invasion factors in this process (T3SS-1, Rck and PagN). Chicks were inoculated with wild-type or isogenic fluorescent *Salmonella* Typhimurium mutants via the intraperitoneal route. Our results show that liver, spleen, gall bladder and aortic vessels could be *foci* of infection and that phagocytic and non-phagocytic cells, including immune, epithelial and endothelial cells, are invaded *in vivo* in each organ. Moreover, a mutant defective for the T3SS-1, Rck and PagN remained able to colonize organs as the wild-type strain and invaded non-phagocytic cells in each organ studied. As the infection of gall bladder was not really described in chicks, invasion of gall bladder cells was confirmed by immunohistochemistry and infection was shown to last several weeks after inoculation of chicks. All together, these findings provide new insights into the dynamics of *Salmonella* spread *in vivo* in chicks at the organ and cellular levels.

## Background

*Salmonella* spp. are among the most important foodborne pathogens. From a public health perspective, according to the World Health Organization, *Salmonella* spp. are among the 31 diarrheal and/or invasive agents (viruses, bacteria, protozoa, helminths, and chemicals) displaying the highest capability of triggering intestinal or systemic diseases in humans. Most cases of salmonellosis are mild, but sometimes the disease is life-threatening and salmonellosis is the third leading cause of death among food-transmitted diseases (1).

The two most commonly reported non-typhoïdal serovars *Salmonella enterica* subsp. *enterica* serovar Enteritidis and *Salmonella enterica* subsp. *enterica* serovar Typhimurium (including its monophasic variant) accounted for almost 80% of human cases occurring in the EU (2). Depending on host factors and serovars, *Salmonella* can induce a wide range of diseases ranging from systemic to asymptomatic infections and gastroenteritis (3). In humans, localized infections can be followed by bacteremia in 3 to 10% of cases (4).

Animals are the primary source of these pathogens and humans become infected mainly by ingesting contaminated food. Poultry meat and eggs are the main source of human *Salmonella* contamination. In 2010, Knight-Jones *et al*. reported that poultry was implicated as an outbreak source in 10.4% of the total cases worldwide (5). Since 2018, it has remained the highest prevalence of *Salmonella*-positive single samples from official control investigations (2). The detection and eradication of *Salmonella* in poultry is difficult because *Salmonella* mostly induce an asymptomatic infection, accompanied by high fecal excretion, which is a source of transmission (6). This leads to contaminated poultry flocks that must therefore be eradicated and all derived products destroyed, resulting in high economic losses. It is therefore particularly important to control animal infection not only to avoid economic consequences but also for the negative impacts for human health.

To establish infection in their hosts, *Salmonella* have to interact with several phagocytic and non-phagocytic eukaryotic cells. Invasion of these cells is considered as one of the most important steps in *Salmonella* pathogenesis. The most well-described invasion process requires the Type III Secretion System – 1 (T3SS-1) encoded by *Salmonella* pathogenicity island 1 (SPI-1). The T3SS-1 is a needle-like structure, which directly injects bacterial effector proteins into the host cytosol to manipulate the cell cytoskeleton, allowing bacterial internalization into non-phagocytic cells (7).Two other *in vitro* entry pathways, involving the Rck and PagN invasins, have also been described in *Salmonella* (8-10). Contrary to the T3SS-1, each invasin interacts with an eukaryotic receptor, EGFR and the heparinated proteoglycan for Rck and PagN, respectively (11, 12). *In vivo*, several reports particularly in mice have shown the key role of the T3SS-1 for *Salmonella* to cross the intestinal barrier (13, 14). Nevertheless, infections in the absence of T3SS-1 in mice, chicks and calves have also been described in several papers in which a mutant, defective for the T3SS-1, was shown to colonize its host as well as its wild-type parent (15-20). This observation has also been made in humans in whom food-borne disease outbreaks have been described with *Salmonella* Senftenberg isolates which lack segments of SPI-1 (21). In the same way, a study performed by Suez *et al*. comparing the pathogenicity of different non-typhoïdal strains concluded that *Salmonella* virulence factors, including multiple T3SS effectors, were absent from several bacteremia isolates suggesting they are dispensable for invasive infection (22). Less is known about the role of the Rck and PagN invasins *in vivo* but *pag*N (formerly *iviVI-A*) and *rck* mutants are both less competitive than their wild-type parent in mice (23-25). However, apart from these roles identified at the organ level, very little is known about the cells targeted by these invasion factors and this is even more true in farm animals.

This topic is crucial because cell and tissue tropism governs disease in many models (26, 27). Moreover, some studies have shown that depending on the entry mechanism both bacterial behavior and host response are different (28). It is therefore important to identify the host cells targeted by *Salmonella* and the different entry routes used by this pathogen to invade the different host cells. In order to improve understanding of how *Salmonella* infect chicks, our aim was to identify cells that could be targeted *in vivo* by this pathogen, expressing or not the known invasion factors. For this purpose, we used a fluorescent *S*. Typhimurium wild-type strain and its fluorescent mutant derivatives deleted of either the T3SS-1 alone or of the three known invasion factors (T3SS-1, Rck and PagN) to infect chicks intraperitoneally. Among the different organs, the spleen, liver and gall bladder were chosen to be potential foci of infection (29). Moreover, vessels from the aortic arch with the brachiocephalic trunk called “aortic vessels” in the article were also collected to analyze putative infection of endothelial cells. In these organs and vessels, identification of phagocytic and non-phagocytic cells and their invasion by the different bacteria were followed using flow cytometric analyses and confocal microscopy.

## Methods

### pFPV-TurboFP650 plasmid construction

Gene encoding TurboFP650 was amplified from the plasmid pTurboFP650-N (Evrogen, Euromedex, France) with primers TurboFP650-XbaI 5’TGCTCTTAGATTTAAGAAGGAGATATAGATATGGGAGAGGATAGCGAGCTG3’ and TurboFP650-SphI 5’CATGCATGCTTAGCTGTGCCCCAGTTTGCTAGG3’. Then, the PCR product and the pFPV25.1 plasmid (66) were restricted by XbaI and SphI restriction enzymes, ligated and transformed into *E. coli* MC1061 (67). pFPV-TurboFP650 recombinant plasmids were selected on Trypticase Soya Agar (TSA - BioMérieux) containing 100 µg/mL of carbenicillin (Sigma-Aldrich) and clones which showed a purple color were selected for restriction analysis. Clones with good restriction profiles were then sequenced to confirm the absence of mutations in the TurboFP650 coding sequence.

### Strains used and inocula preparation

The pFPV-TurboFP650 plasmid was introduced in *S*. Typhimurium 14028 wild-type (WT), the Δ*invA*::*kan* mutant (Δ*invA*; T3SS-1 defective) or the Δ*invA::kan* Δ*pagN::cm* Δ*rck* mutant (3Δ) (30).

To prepare the inocula, the strains were cultured in Trypticase Soya Broth (TSB - BioMérieux) supplemented with carbenicillin 100 µg/mL for 24h at 37°C with shaking. The cultures were centrifuged at 4500g for 20 min at 20°C and the pellets were suspended in phosphate buffered saline (PBS) containing 50% glycerol. The bacterial suspensions were then aliquoted, frozen and stored at −80°C. The frozen aliquots from the same initial inoculum were used throughout the experiments.

### Experimental infection

Five-day-old PA12 White Leghorn chicks, provided by the Experimental Platform for Infectious Disease (UE 1277 - INRAE) were intraperitoneally inoculated with 0.2 mL of bacterial suspension. On the day of inoculation, a frozen aliquot of the inoculum was thawed. Bacteria concentrations were standardized turbidimetrically and diluted to a concentration of 6.107 CFU/0.2 mL in PBS. Chicks were maintained in medium isolator systems (0.83 m^2^) with controlled environmental conditions (feed, water, temperature, air humidity and lighting scheme) for two days before sacrifice by decapitation and bleeding. To follow the persistence in the gall bladder, the inoculation dose was 3.10^7^ CFU/chick, in order to reduce the mortality of chicks observed with the higher dose. The kinetics of organ colonization was followed each week over a period of 36 days.

### Enumeration of bacterial load in infected organs

On the day of sacrifice, control animals of the same age (i.e. not inoculated) were provided by the Experimental Platform for Infectious Disease. Spleens, livers, gall bladders and the aortic vessels were collected aseptically from each animal for quantification of bacterial load.

To determine the bacterial load, organs were homogenized in TSB and serial 10-fold dilutions were plated on TSA or *Salmonella*–*Shigella* medium supplemented with carbenicillin 100 µg/mL. The colonies per plate were counted after incubation for 24 h at 37°C. Counts were expressed as log (CFU) per g of organ.

### Preparation of cells for flow cytometry

For the flow cytometric analyses, organ-specific samples were obtained by pooling the spleens, livers, aortic vessels and gall bladders of the different chicks in Hanks’ buffered saline solution (HBSS) without Ca^2+^ and Mg^2+^ in the dark at 4°C in order to be able to analyze at least 200,000 cells for each organ. Independent infections were repeated at less three times. Gall bladders and aortic vessels were cut into small pieces and samples put in collagenase A (0.3% - Sigma) – dispase I (1 U/mL - Sigma) – HBSS for 30 min at 37°C. The whole purification process was performed at 4°C. All organs were then homogenized in HBSS using syringe plungers and filtered through 40-μm-mesh cell strainers (Falcon), before being transferred into a 50-ml centrifuge. After centrifugation at 1000*g* for 15 min, cells were washed, resuspended in HBSS at approximatively 5.10^6^ – 1.10^7^ cells / mL and maintained in the dark at 4°C.

### Flow cytometric analyses

Cells were characterized according to the antibodies available in poultry (S2 Table). Mouse Anti-Chicken antibody, clone KUL01 specifically recognizes chicken monocytes, macrophages and interdigitating cells (68). Anti-CT3 antibody targets the avian homolog of the CD3-antigen, a common antigen used to identify T lymphocytes (69). Clone AV20 antibody recognizes the antigen Bu-1, a chicken B-cell marker, commonly used to identify B lymphocytes (70). Mouse anti-chicken CD41/61 clone 11C3 recognizes chicken integrin CD41/61 that is expressed on chicken thrombocytes and cells of the thrombocyte lineage (71). Mouse anti L-CAM antibody recognizes an 81 kDa N-terminal tryptic fragment of L-CAM, an epithelial cell marker, from embryonic chicken liver plasma membranes (72) and last VE-Cadherin is an intercellular junction marker of endothelial cells. This is a synthetic peptide corresponding to Human VE-Cadherin amino acids from position 750 to the C-terminus conjugated to keyhole limpet hemocyanin. Rabbit polyclonal antibody anti-VE-cadherin clone reacts with mouse, chicken and human VE-Cadherin.

The anti-Bu-1 and the anti-CD3, that allow B and T lymphocytes to be identified, were FITC conjugated. Antibodies that allow identification of monocytes-macrophages, thrombocytes, epithelial and endothelial cells, required an Alexa Fluor™488 conjugated anti-secondary anti-mouse or anti-rabbit antibodies. The endothelial cell samples were pre-treated with 20% horse serum. The primary antibodies were incubated with cells for 90 min at 4°C in the dark and then rinsed in HBSS. When necessary, secondary antibodies were added for 90 min at 4°C in the dark, and then rinsed. Appropriate isotype control antibodies (S2 Table) were used to determine the levels of unspecific staining in all the experiments. Parallel samples were stained with a Fixable Viability Dye Cell Staining eFluor 450 (eBioscience (65-0863)) to determine the settings for a live cell gate based on light scatter properties. All samples were then filtered through 60-µm nylon Blutex just before flow cytometric analyses were performed using a BD LSR Fortessa™X-20 (BD Biosciences, San Jose, CA, USA). BD FACSDiva™software (v 8.0.2, RRID:SCR_001456) was used to analyze the cytometric data. Infected and control samples were manipulated under the same conditions.

### Identification and relative quantification of infected and non-infected cells by flow cytometry

For each sample, dot plots were analyzed. The intensity of green fluorescence (FITC or Alexa Fluor™ 488) is on the vertical axis, plotted against the intensity of red fluorescence (TurboFP650) on the horizontal axis. Labelled infected cells thus emitted both green and red fluorescence. They were revealed as dots in the upper right-hand part of the graph. For each experiment in each organ, a gate was determined removing inappropriate labeling - debris based on morphological criteria. Regions were set according to uninfected control samples and isotype-control staining. In order to have quantitative results, 200,000 events were analyzed for each sample for all staining. Examples are provided for some cell-type/organ labeling (for monocytes-macrophages (S1 Fig) and thrombocytes (S2 Fig) in the gall bladder, B lymphocytes (S3 Fig) and T lymphocytes (S4 Fig) in the spleen, epithelial cells in the liver (S5 Fig) and endothelial cells in the aortic vessels (S6 Fig)). Quantification of the percent of positively labeled cells was then calculated by subtracting the number of cells in the control areas from those in the positive labeled areas. The positive labeling areas of B and T lymphocytes cells were established using a control mouse IgG1-FITC conjugate, whereas the positive labeling areas of monocytes-macrophages, thrombocytes and epithelial cells were determined with a control mouse IgG1-Alexa Fluor™ 488 conjugate. For the labeling of the endothelial cells, we used rabbit IgG, followed by a secondary antibody, an anti-IgG-Alexa Fluor™488 conjugate. The total percentages of infected cells, positive labeled or unlabeled were also determined. All negative responses were scored at 0.001% to account for the threshold and allow for a logarithmic representation of the results. The medians are represented by a red dash.

### Purification of the infected cells and confocal laser-scanning analysis

Cells were sorted using a high-speed cell sorter, MoFlo Astrios ^EQ^ (Beckman Coulter Inc, Brea, CA, USA) equipped with four lasers: violet (405nm), blue (488nm), yellow-green (561nm) and red (640nm) and placed under a class II biological safety cabinet. We used a nozzle of 90 µm and selected a sheath pressure of 40 psi. Sorted cells were collected in 1.5 ml Eppendorf tubes containing 350 µL of HBSS medium supplemented with 10% fetal calf serum to limit cell stress.

After cell sorting, samples were deposited on glass coverslips and centrifuged with a cytospin at 200 rpm for 10 min. Cells were then fixed in formaldehyde 4% for 10 min. Nucleus staining was performed with DAPI 1 µg/mL for 1 min and coverslips were mounted on slides with fluorescent mounting medium (Dako). Cells were observed under a SP8 confocal laser-scanning microscope equipped with an HCP PL APO 100x/1.44 Oil CORR CS immersion objective (Leica). Z-stacks were re-sliced horizontally and vertically to obtain the projections of perpendicular views from 3D images, providing a view of all bacteria in the cells, using Las AF lite 2.6.3 build 8173 software (Leica application Suite X, RRID:SCR_013673).

### Immuno-histochemistry (IHC)

Chick gall bladders were fixed in 4% buffered paraformaldehyde at 4°C for 24 h. They were then processed by routine methods, paraffin embedded, cut in sections (thickness, 5 µm), and stained for IHC with HRP detection. All samples were incubated at room temperature. The primary antibody was an anti-*Salmonella* lipopolysaccharide marker: rabbit anti-*Salmonella* O:4,5 (1/100 – D. Pasteur). The tissue sections were dewaxed in Histosol (Shandon, Thermoscientific), rehydrated in a decreasing series of ethanol, rinsed and rehydrated in tap water. Sections were treated with heat-induced epitope retrieval, 10mM Sodium Citrate buffer, pH 6, 121°C, for 15 min. The tissues were then rinsed in tap water. The endogen peroxidase was blocked in 1% hydrogen peroxide and methanol for 30 min. Preparations were rinsed in PBS with 1% skimmed milk and 0.05%Tween 20 (PBSTM), blocked in 20% goat serum - 30% fetal calf serum - PBS for 20 min. They were then incubated with a primary antibody for 60 min and rinsed in PBSTM, followed by N-Histofine rabbit, HRP (MMFRANCE) for 30 min. At the end, samples were rinsed in PBSTM, incubated with chromogen (diaminobenzidine, liquid DAB +, MMFrance) for 5 min, counterstained with hematoxylin of Harris (Merck, Labellians), rinsed in tap water, dehydrated in successive ethanol baths (50°, 70°, 90° and absolute, each for 2 min), cleared in histosol, and mounted on coverslips with Eukitt.

Tissues were examined and photographed with a light microscope Eclipse 80i, Nikon with DXM 1200C digital camera (Nikon Instruments, Europe, Amsterdam, Netherlands) and NIS-Elements D Microscope Imaging Software (NIS-Elements, RRID:SCR_014329).

### Statistical analyses

A Kruskal-Wallis Test was conducted to examine the differences in the levels of organ colonization, followed by a Dunn’s multiple comparisons test (GraphPad Prism version 6.07 for Windows, GraphPad, www.graphpad.com, RRID:SCR_002798). Significance was *p < 0.05 and **p < 0.01.

For the flow cytometric analyses, asymptotic two-sample Fisher-Pitman permutation tests (One-Way-Test) were performed with R software, package Rcmdr version 2.5.3 (R Project for Statistical Computing, RRID:SCR_001905). Significance was * p<0.05 (http://www.r-project.org, http://socserv.socsci.mcmaster.ca/jfox/Misc/Rcmdr/).

## Results – Discussion

### Mutant strains defective for the T3SS-1 or the three known invasion factors inoculated by the intraperitoneal route, colonize chicks more effectively than their wild-type parent strain

Previous work has shown that a Δ*invA* mutant strain (T3SS-1 defective strain) and a strain deleted for the three known invasion factors (3Δ) remained invasive for several eukaryotic cell lines compared to the wild-type strain (30). To determine the ability of our wild-type strain and our mutant strains to invade *in vivo* several host cells, we chose to bypass the intestinal barrier and to infect chicks by the intraperitoneal route. The first step was to evaluate the ability of these different strains to colonize vessels and several organs of chicks i.e. the spleen, liver and gall bladder (Fig 1). To ensure that the levels of bacteria were not related to the presence of *Salmonella* in the blood, all chicks were bled.

**Fig 1.**
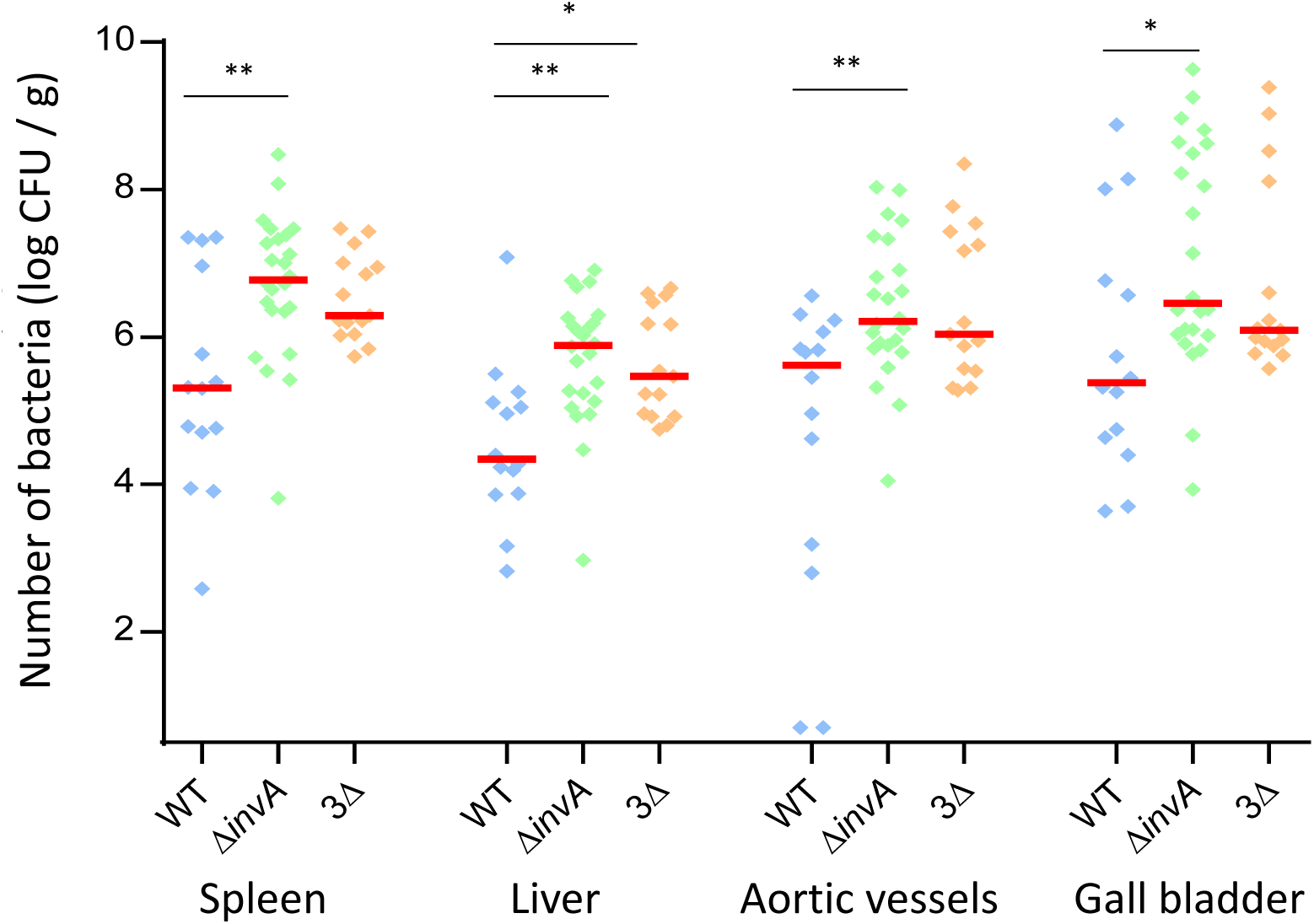
Level of different *S*. Typhimurium strains in organs of chicks after intra-peritoneal inoculation. Five-day-old chicks were intraperitoneally inoculated with around 6.10^7^ CFU/chick with *S*. Typhimurium 14028 turboFP650 wild-type strain (WT 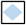), Δ*invA*::*kan* mutant strain (Δ*invA*; T3SS-1 defective 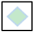) or the *ΔinvA::kan ΔpagN::cm Δrck* mutant strain (3Δ; T3SS-1, Rck, PagN defective 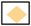). Two days post-infection, spleens, livers, aortic vessels and gall bladders were removed aseptically from each animal for quantification of bacterial load. Results are expressed as number of bacteria per g of organ (log CFU per g of organ). The medians are represented by a red dash. A Kruskal-Wallis Test was conducted, followed by Dunn’s multiple comparisons test (GraphPad Software). Significance was *p < 0.05 and **p < 0.01.

The first observation is that all strains were able to infect all the organs and vessels. For some animals, the infection rate even reached 8 log CFU/g, especially for the gall bladder. Moreover, all organ colonization levels were higher after inoculation with the Δ*invA* strain compared to that with the wild-type strain (29, 35, 3.9 and 12 times more in the spleen, liver, aortic vessels, and gall bladder, respectively). All these differences were statistically significant. While the 3Δ strain also colonized these organs and vessels, more effectively than the wild-type strain (9.5, 13, 2.6 and 5.1 times more in the spleen, liver, aortic vessels, and gall bladder, respectively), a statistically significant difference was only identified in the liver. No significant statistical difference could be observed between the two mutants, beside the fact that the log CFU of bacteria/g for the 3Δ strain was always inferior to that of bacteria/g of organ for the Δ*invA* strain. The levels of CFU recovered in the gall bladder and the aortic vessels should be highlighted, as they have previously been rarely studied. These results demonstrated that all the tested organs and vessels are colonized by *S*. Typhimurium and that the two mutant strains (Δ*invA* or 3Δ) colonized deep organs of chicks after intraperitoneal inoculation at least at the same level as the wild-type strain.

This latter result could be attributed to the route of inoculation. Indeed, *Salmonella* injected *via* the intraperitoneal route easily reaches systemic sites such as the spleen and the liver. They could also reach the gall bladder through the vasculature or the hepatic duct (31). *In vivo* studies have demonstrated that the T3SS-1 is primarily associated with the early stage of infection in which it translocates T3SS1 effectors across the host intestinal epithelial cell membrane and stimulates intestinal inflammation (13) (14, 32-34) and is therefore important for *Salmonella* colonization after animals are inoculated orally. Our results show that, in chicken, the colonization of deeper organs can be independent of this type III secretion system as no difference in colonization between a T3SS-1 mutant and its wild-type parent was observed as described after intraperitoneal or intravenous inoculation of mice. Several reports have also suggested that *Salmonella* remain pathogenic without an active T3SS-1 even after oral inoculation in several animal models, including chicks (19, 20, 35). Moreover, a *Salmonella* Senftenberg strain lacking T3SS-1 was isolated from a human clinical case and has been shown to be able to induce enterocolitis in a mouse model (21). In our case, one hypothesis that could explain the higher colonization of the mutant strains compared to the wild-type strain is that, after intraperitoneal inoculation, the absence of the T3SS-1 could induce a lower immune system alert, especially a lower inflammatory response and consequently less killing of bacterial. Indeed, SPI-1 genes are involved in the regulation of the host immune response, for example the host inflammatory response (36), immune cell recruitment (37) and apoptosis (38, 39). Moreover, we already know that a SPI-1 mutant and also a *phoP* mutant, not expressing PagN like our 3Δ mutant, did not stimulate an inflammatory response in the caecum of chicks (40).

Cytometric analyzes and microscopy were then performed in order to determine whether *Salmonella* was within the cells of the different organs and vessels and to identify the cell-types infected. The infectious dose of 6.10^7^ CFU/chick used for the previous *in vivo* experiment, represented a good compromise between the infectious dose and the period of slaughter (2 days), in order to potentially detect enough intracellular bacteria for flow cytometry analyses. The animals were bled to decrease red blood cells and allow better detection of organ cells. The concentration of *S*. Typhimurium-TurboFP650-wild-type strain was checked in the blood of six animals. An average of 1.95 ± 1.09 log CFU/mL was found.

### STM-Turbo FP650-WT and its mutant strains were within the cells and did not only adhered to the cells

As our aim was to identify cells infected by *Salmonella*, we first assessed whether our protocol allowed us to identify intracellular bacteria or not. Indeed, flow cytometry is useful for quantitative analyses but it does not allow the intracellular localization of bacteria to be determined as adherent bacteria could exist. According to our protocol, it was highly unlikely that *Salmonella* would only be present extracellularly due to the methods used to purify and mechanically separate the cells, including filtrations and washings and, for some organs, enzymatic cleavage with two different enzymes (collagenase and dispase) were performed for cell purification. Theoretically, after all these treatments related to organ dissociation, only a few bacteria would remain adhered, suggesting that the large majority were intracellular. However, in order to confirm this, cell sorting based on the labeling of the cells and confocal analyzes were carried out for each cell type of each organ. For flow cytometry, regions corresponding to infected cells were identified with the PE-cy5 canal and were set according to uninfected control samples. The Alexa fluor 488 canal was used to identify the cell types according to the isotype-control staining. One example for each cell type is given in Fig S1, Fig S2, Fig S3, Fig S4, Fig S5 and Fig S6. Double-labeled cells were sorted using flow cytometry and observed with confocal microscopy. A Z-stack was re-sliced horizontally and vertically to obtain the projections of perpendicular views, confirming the intracellular presence of bacteria. This allowed us to observe intracellular *Salmonella* expressing red tag, in green labeled cells for all the cells considered. These results confirm the intracellular localization of the different strains and thus validate our protocol designed to identify and quantify the cell types infected by *Salmonella* in selected chick organs. Moreover, they show that *S*. Typhimurium can invade all the cell types studied in this paper, i.e. monocytes-macrophages, B and T lymphocytes, thrombocytes, epithelial and endothelial cells of chicks. Currently, only a few papers have described the cells infected by *Salmonella in vivo* and most of these papers are in mouse models. In these articles, *Salmonella* were found mainly in macrophages and neutrophils from the liver and spleen of mice, but infected B and T lymphocytes were also identified (41-44).

A more detailed analysis of the confocal images allowed us to observe that in most cases infected cells, whatever the cell type, only harbored one to five bacteria per cell (Fig 2), but in a few, more bacteria were visualized. This result is consistent with results obtained in the literature on *Salmonella* infected macrophages *in vivo*. Indeed, in many experiments in mice, the majority of liver or spleen infected phagocytes contained relatively few bacteria (41-43, 45), but the presence of many bacteria per cells has also been reported (42, 45, 46). Our results show that this heterogeneous number of bacteria per cell could be enlarged to non-phagocytic cells in chicks. However, in mice it seems that the number of bacteria per cell had a moderate impact on the infectious process as host cells that contain high numbers of bacteria have the same probability of undergoing lysis as cells containing only a few bacteria (47). Both highly and weakly infected cells contributed significantly to the *Salmonella* infection process and not only the macrophages (43).

**Fig 2.**
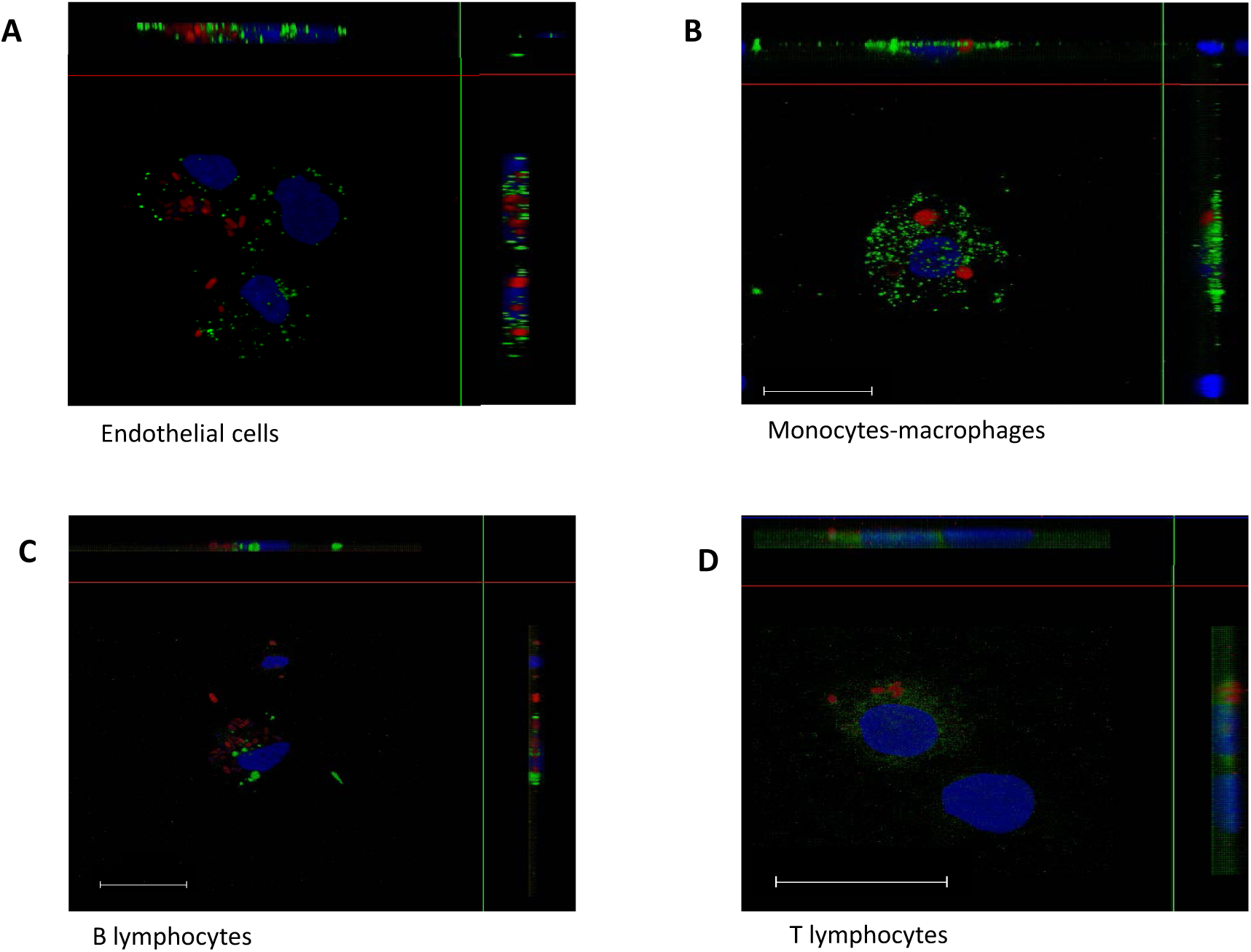
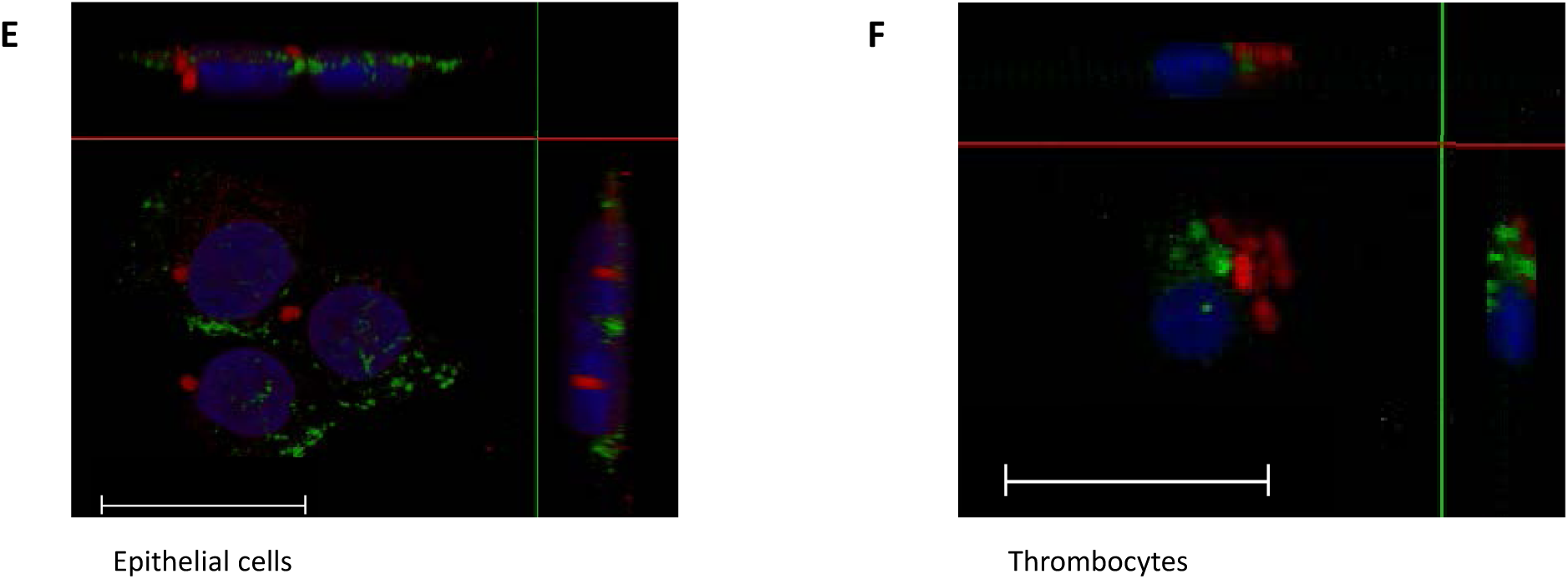
Intracellular localization of *Salmonella* in cells purified from *in vivo* infected organs. Five-day-old chicks were intraperitoneally inoculated with around 6.10^7^ CFU/chick with *S*. Typhimurium 14028 turboFP650 wild-type strain (WT 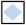), Δ*invA*::*kan* mutant strain (Δ*invA*; T3SS-1 defective 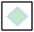) or the *ΔinvA::kan ΔpagN::cm Δrck* mutant strain (3Δ; T3SS-1, Rck, PagN defective 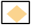). Two days post infection, animals were sacrificed and the different organs removed. After, cells were isolated from organs, they were sorted using a high-speed cell sorter, MoFlo Astrios EQ and deposited on glass coverslips after cytospin at 200 rpm for 10 min. Cells were then fixed in formaldehyde. Nucleus staining was performed with Dapi (blue). The bacteria are in red (turboFP650), whereas cells are identified in green thanks to FITC or Alexa Fluor™488 conjugated antibodies. Cells were observed under a SP8 confocal laser-scanning microscope equipped with a 100x oil immersion objective (Leica). Z-stacks were re-sliced horizontally and vertically to obtain the projections of perpendicular views from 3D images, allowing a view of all bacteria in the cells, using Las AF lite 2.6.3 build 8173 software (Leica). White dashes represent 20 µm. **A** represents endothelial cells from the aortic vessels, infected with the 3Δ strain. Picture size 32.54 µm x 38.45 µm. **B** represents monocytes-macrophages from the liver, infected with the Δ*invA* strain. Picture size 116.25 µm x 116.25 µm. **C** represents B lymphocytes, infected with the wild-type strain. Picture size 58.13 µm x 58.13 µm. **D** represents T lymphocytes, infected with the wild-type strain. Picture size 39.88 µm x 39.88 µm. **E** represents epithelial cells in the gall bladder, infected with the 3Δ strain. Picture size 37.80 µm x 37.80 µm. **F** represents thrombocytes in the aortic vessels, infected with the 3Δ strain. Picture size 116.25 µm x116.25 µm.

### Analysis of the infected cell types in the spleen

In the chicken spleen, the distinction between the red and white pulp is less marked than in mammals. The red pulp mainly contains erythrocytes, granulocytes, macrophages, scattered T lymphocytes and plasma cells. However, the architecture of the avian white pulp differs considerably. Three morphologically distinct areas constitute the spleen. The first consists of peri-arteriolar lymphocyte sheaths, mainly containing T lymphocytes that surround arterioles, which have visible muscular layers. The second involves peri-ellipsoid lymphocyte sheaths (PELS) surrounding capillaries, lacking muscular tissue and lined by cuboidal endothelium and reticulin fibers. The last consists of follicles with germinal centers, surrounded by a capsule of connective tissue. PELS and follicles mainly contain B lymphocytes (48).

In the spleen, the six antibodies used in our study allowed us to detect about 86% of the total cells. Epithelial cell labeling was not necessary, as these cells were not expected to be present. B lymphocytes (average of 8%), T lymphocytes (average of 22%) and endothelial cells (average of 30%) were identified the most, as expected (Fig 3A). Compared to the non-infected chicks, the percentage of labeled cells was similar in the groups of chicks inoculated with the wild-type bacteria, the single or triple mutant bacteria. The only statistical difference was observed for the percentage of thrombocytes between the uninfected chicks and the chicks infected with the wild-type strain (p=0.049. S1 Table). The small decrease in the number of thrombocytes after infection with the wild-type strain, was similar to that observed with the two mutants, but the number of independent experiments was probably not sufficient to obtain a statistical difference between the uninfected group and the chicks inoculated with these *Salmonella* mutants. Similarly, the lower percentage of macrophages observed after infection with the 3Δ mutant strain was not significant. Either the absence of cell recruitment is real in chicks or it could be related more to the fact that the infected cells were identified only two days after the intra-peritoneal inoculation. The infection rates observed for all the identified cells (lymphocytes, macrophages, thrombocytes, and endothelial cells) were between 0.1 and 1%. Monocytes and macrophages were proportionally the most infected cells (about five times more than the other cell types), but endothelial cells, and B and T lymphocyte cells were the most infected cells in the spleen as their absolute number was higher than that of monocytes-macrophages in this organ. No statistical differences were observed according to the *Salmonella* strains tested (Fig 3B). The fact that monocytes-macrophages were identified as being proportionally the most infected cells of the spleen was not surprising. In mice, it is commonly assumed that the systemic spread of *Salmonella* is contingent upon dissemination and survival within macrophages. Indeed, survival in macrophages is essential for virulence (49). However, contrary to what was assumed, our work, clearly demonstrated that other cell types, such as lymphocytes, thrombocytes and endothelial cells could also be infected by *Salmonella* in chicken spleen. As monocytes and macrophages are phagocytic cells, the fact that there was no difference in the percentage of monocyte-macrophage infected cells between the mutants and the wild-type strain was to be expected. By contrast, B and T lymphocytes, thrombocytes and endothelial cells are non-phagocytic cells and thus a difference in the percentage of cells infected by the different strains could have been expected. However, Geddes *et al*. have also described in mice the internalization of *Salmonella* in splenic B and T cells, independently of the T3SS-1 (44). Our work suggests that this observation could be extended to other non-phagocytic cells of other animal species.

**Fig 3.**
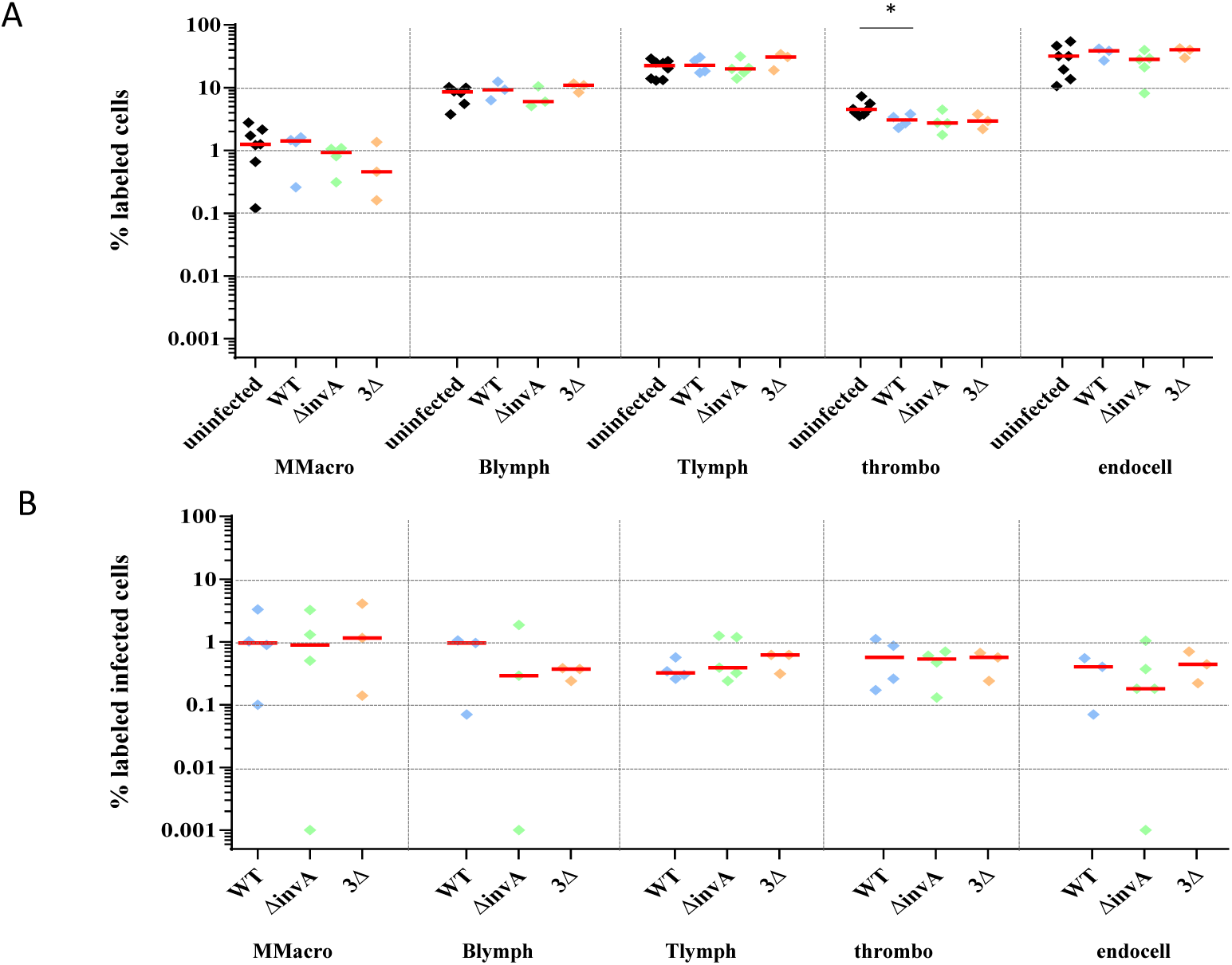
Percentage of identified and *Salmonella* infected cells in spleen. Five-day-old chicks were intraperitoneally inoculated with around 6.10^7^ CFU/chick with *S*. Typhimurium 14028 turboFP650 wild-type strain (WT 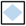), Δ*invA*::*kan* mutant strain (Δ*invA*; T3SS-1 defective 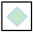) or the *ΔinvA::kan ΔpagN::cm Δrck* mutant strain (3Δ; T3SS-1, Rck, PagN defective 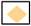). Two days post infection, animals were sacrificed and the different organs removed. Cells from uninfected animals of the same age were used as a control. After labeling with the corresponding antibodies, the percentages of macrophages-monocytes, B and T lymphocytes, thrombocytes and epithelial and endothelial cells were quantified by flow-cytometry. The percentage of labeled cells (A) and the percentage of labeled infected cells (B) are represented. All negative responses were scored at 0.001%. The medians are represented by a red dash. Asymptotic two-sample Fisher-Pitman permutation tests (One-Way-Test) were performed (R software). Significance was * p<0.05.

### Analysis of the infected cell types in the liver

The liver is divided into a right and a left lobe. Each lobe of the liver has approximately 100,000 lobules separated from each other by interlobular *septum*. These lobules are formed by parenchymal cells (hepatocytes), which represent 80% of the total liver volume and non-parenchymal cells localized in the sinusoidal wall. These sinusoidal walls are the vascular side of the hepatocytes and they are composed of endothelial cells and macrophages. These macrophages are star-shaped and confined to the liver. Called Kupffer cells, they phagocyte pathogens, cell debris and damaged red and white blood cells (50).

In the liver, the six antibodies used allowed us to detect about 76% of the cells and we detected as many epithelial cells (average of 31%) as endothelial cells (average of 33%) (Fig 4A). As expected, these were the main cell types identified. Few monocytes-macrophages were identified. One hypothesis is that the KUL01 antibody poorly recognizes the Kupffer cells (51). Another hypothesis is that their percentage compared to epithelial and endothelial cells is very low in the liver. There were also very few, if any, T lymphocytes. In humans and mice, lymphocytes are present in small quantities at the level of the sinusoids and the space of Disse (perisinusoidal space) and histological investigation does not suggest that there are many immunologically relevant cells present (52). Liver-resident lymphocytes serve as sentinels and perform immunosurveillance in response to infection and non-infectious insults, and are involved in the maintenance of liver homeostasis (53). Our low level of T lymphocytes in the liver is most probably related to the fact that our observations were made two days after the intraperitoneal inoculation and that our chicks were only six days old and therefore immunologically immature. For all cell types, in the liver, the percentage of labeled cells was quite similar, whatever the infected or uninfected status of the animals. Only a statistical difference for the percentages of labeled epithelial cells between the uninfected chicks and those infected with the *invA* mutant was observed (p=0.039. S1 Table). Like in the spleen, we were able to observe similar levels of infected cells between chicks inoculated with the wild-type strain or with the two mutant strains deleted of the known entry factors. Compared to the spleen, the percentages of labeled infected cells were more heterogeneous (Fig 4B). In particular, the percentages of infected monocytes-macrophages and B lymphocytes were around 3%, while the percentages of infected epithelial and endothelial cells were 0.10% and 0.24%, respectively. Nevertheless, as these latter cell types are more frequent in the liver than monocytes-macrophages and B lymphocytes (Fig 4A), endothelial and epithelial cells represent a large proportion of the infected cells in the liver. The monocytes-macrophages of the liver are responsible among others, for the phagocytosis of pathogens and thus, it is not surprising that they were found infected, with no differences whatever the strain inoculated. In contrast, the thrombocytes were very weakly infected here.

**Fig 4.**
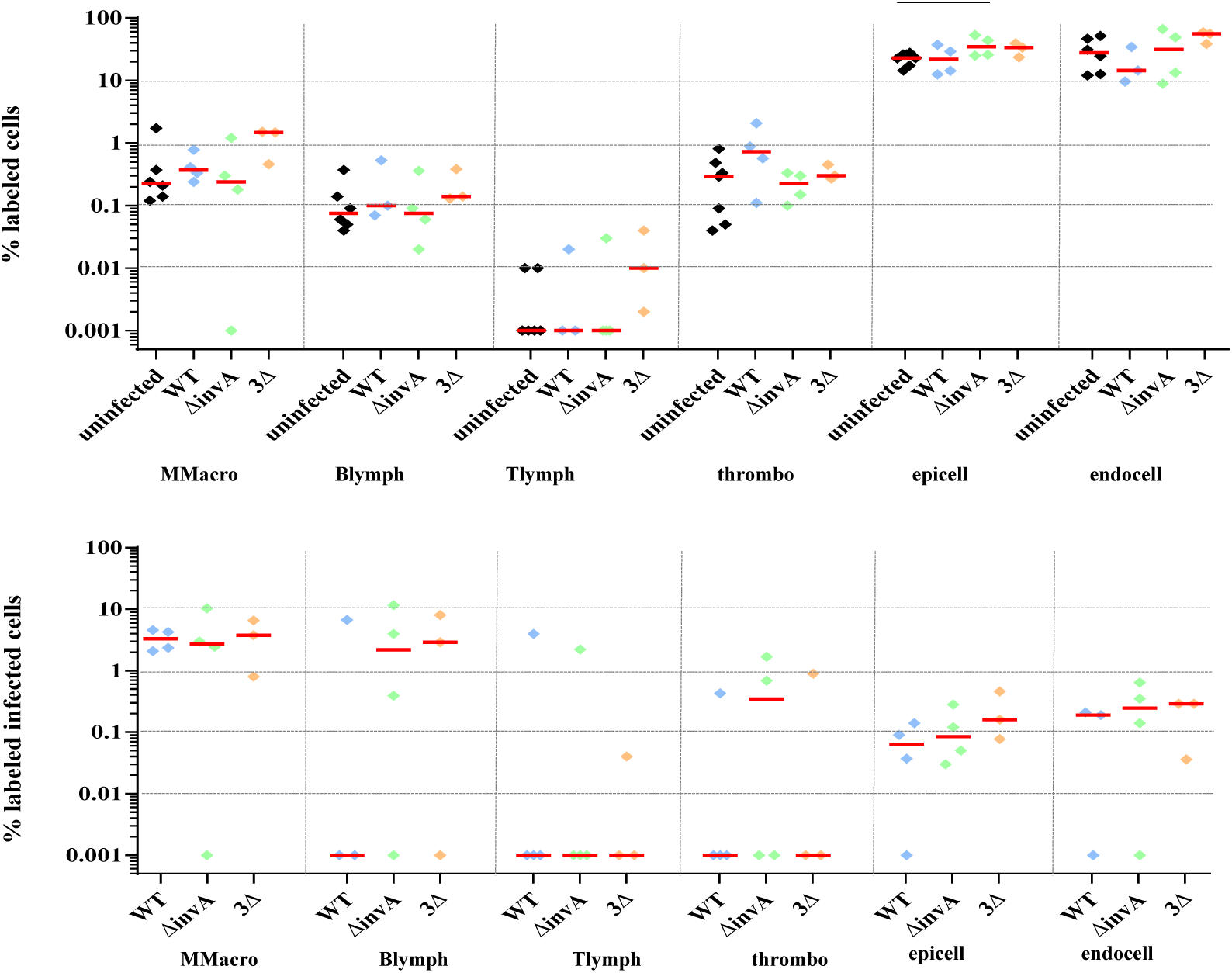
Percentage of identified and *Salmonella* infected cells in liver. Five-day-old chicks were intraperitoneally inoculated with around 6.10^7^ CFU/chick with *S*. Typhimurium 14028 turboFP650 wild-type strain (WT 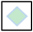), Δ*invA*::*kan* mutant strain (Δ*invA*; T3SS-1 defective 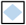) or the *ΔinvA::kan ΔpagN::cm Δrck* mutant strain (3Δ; T3SS-1, Rck, PagN defective 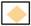). Two days post infection, animals were sacrificed and the different organs removed. Cells from uninfected animals of the same age were used as a control. After labeling with the corresponding antibodies, the percentages of macrophages-monocytes, B and T lymphocytes, thrombocytes and epithelial and endothelial cells were quantified by flow-cytometry. The percentage of labeled cells (A) and the percentage of labeled infected cells (B) are represented. All negative responses were scored at 0.001%. The medians are represented by a red dash. Asymptotic two-sample Fisher-Pitman permutation tests (One-Way-Test) were performed (R software). Significance was * p<0.05.

All together, these results strengthen those observed in the spleen, showing that at least four different cell types, i.e. monocytes-macrophages, B lymphocytes, endothelial and epithelial cells, were infected by *Salmonella* in the liver of chicks.

### Analysis of the infected cell types in the aortic vessels

The term “aortic vessels” in our paper corresponds in fact to the aortic arch and the brachiocephalic trunk. In contrast to mammals, two brachiocephalic trunks arise from the arch of the aorta and give rise to the common carotid and subclavian arteries in birds (54). This “organ” was chosen as a source of endothelial cells.

For this organ, only 55% of the cells were identified through flow cytometric analysis. This low percentage of identified cells was mainly related to the presence of smooth muscle fibers in vessels for which no any antibodies exist for the chicken. The *adventitia*, which is the outer layer of the arterial wall, is made up of connective tissue and elastic fibers. It contains capillary vessels vascularizing the arterial wall as well as nerve fibers of the sympathetic and parasympathetic autonomic system. According to the size of the arteries, the media, which is the middle layer of the arterial wall, is made up of collagen, elastin or smooth muscle fibers allowing vasoconstriction. The *intima*, the inner layer of the arterial wall separated from the media by the internal elastic limiter, is formed of the vascular endothelium (cell monolayer) resting on a layer of connective tissue (55).

The percentages of labeled cells in aortic vessels according to the chick group illustrated in Fig 5A were more dispersed than for the previous two organs, probably due to the treatment of the aortic vessel with enzymes, which made extraction less easy. Monocytes-macrophages and endothelial cells were the most representative type of cells labeled, but all cell types were identified (Fig 5A). This is the first organ in which we could observe a difference between the percentages of monocytes-macrophages according to the uninfected or infected status of the animals. Contrary to the spleen and the liver, the percentage of labeled monocytes-macrophages showed statistically significant differences between the uninfected chicks and those infected with the wild-type strain, on one hand, (p = 0.035. S1 Table) and those infected with the 3Δ mutant, on the other hand (p = 0.043. S1 Table). Despite a high percentage of labeled endothelial cells, few if any were infected. By contrast, all the other cell types were infected and to a greater proportion than in the spleen and liver (Fig 5B). Indeed, compared to the spleen and liver, the median percentage of each infected cell type in the aortic vessels, except endothelial cells, varied from 1 to 10% versus 0.1 – 1% in the spleen or 0 to 3% in the liver. Surprisingly, in some chicks more than 10% of lymphocytes, thrombocytes and epithelial cells were infected.

**Fig 5.**
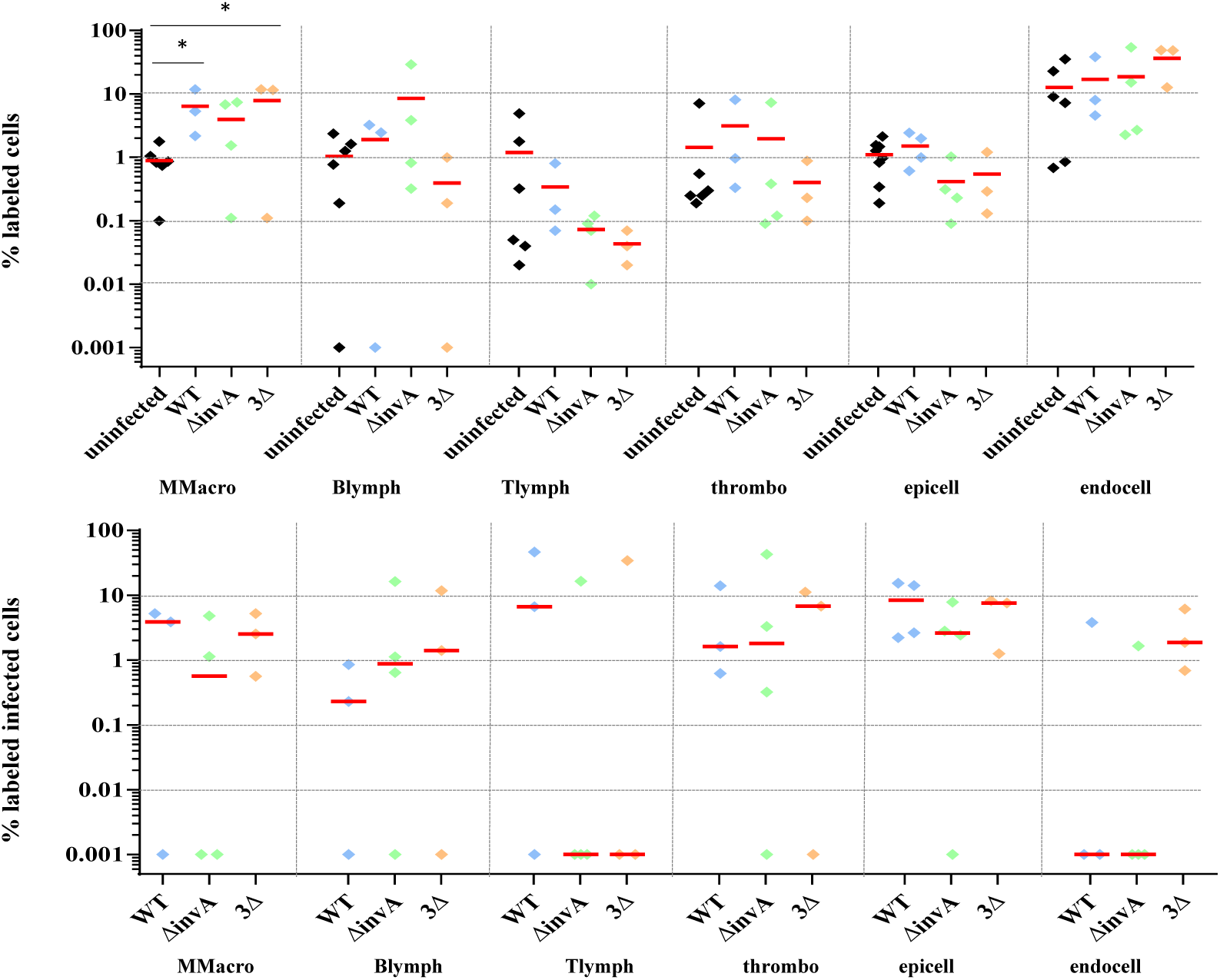
Percentage of identified and *Salmonella* infected cells in aortic vessels. Five-day-old chicks were intraperitoneally inoculated with around 6.10^7^ CFU/chick with *S*. Typhimurium 14028 turboFP650 wild-type strain (WT 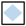), Δ*invA*::*kan* mutant strain (Δ*invA*; T3SS-1 defective 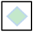) or the *ΔinvA::kan ΔpagN::cm Δrck* mutant strain (3Δ; T3SS-1, Rck, PagN defective 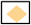). Two days post infection, animals were sacrificed and the different organs removed. Cells from uninfected animals of the same age were used as a control. After labeling with the corresponding antibodies, the percentages of macrophages-monocytes, B and T lymphocytes, thrombocytes and epithelial and endothelial cells were quantified by flow-cytometry. The percentage of labeled cells (A) and the percentage of labeled infected cells (B) are represented. All negative responses were scored at 0.001%. The medians are represented by a red dash. Asymptotic two-sample Fisher-Pitman permutation tests (One-Way-Test) were performed (R software). Significance was * p<0.05.

These results clearly show that, like in the spleen and liver, numerous cell types are infected in vessels.

### Analysis of the infected cell types in the gall bladder

The avian gall bladder is attached to the right liver lobe. Histologically, the avian gall bladder is composed of three *tunicae*. The first, the *tunica mucosa* is mainly lined with non-ciliated simple columnar epithelium and consists of a layer of connective tissue with elastic and muscle fibers. The second, the *tunica muscularis* consists of smooth muscle fibers and abundant intervening connective tissue. The third, the *tunica serosa* consists of coarse collagen fiber and elastic fibers. All epithelial cells are basally located and contain an oval nucleus. Bile is synthesized in the hepatocytes and secreted into bile canaliculi located on the lateral surfaces of adjoining liver cells (56). Relatively little is known about biliary secretion in birds due to the complex anatomy in which bile enters the intestine via both hepato-enteric and cystico-enteric ducts. In ruminants, pigs and poultry, there is relatively continuous secretion of bile into the intestine (50).

About 75% of cells were identified with the available antibodies. Numerous “unidentified cells” would most probably correspond to fibroblasts. Endothelial cell labeling was not performed, as they were not expected to be found in the gall bladder. By contrast, all other cell types were identified. The epithelial cells represented about 60% of the identified cells and the percentages of thrombocytes and monocytes-macrophages were between 1 and 10% (Fig 6A). B and T lymphocytes were less present. As for the aortic vessels, the variability between animals was considerable, certainly due to the breakdown of organs with different enzymes, which made extraction less reproducible. As in the other organs (except for the percentages of monocytes-macrophages in the aortic vessels), there were no differences in the percentages of the labeled cells between uninfected and infected chicks. By comparing the inoculated chicks, only one statistically significant difference was observed for the percentage of T lymphocytes between the chicks inoculated with the single or the triple mutant (p = 0.041. S1 Table). Immune cells were highly infected. For example, about 10% of the B and T lymphocytes were infected by the different strains. Moreover, B and T lymphocytes in the gall bladder were found to be more infected than in the other organs. The other identified cells were infected between 1 and 10 % (Fig 6B). In all cases, except one, no statistical differences were observed between the chicks inoculated with the wild-type or the mutant strains. The only significant statistical difference was observed for the infected monocytes-macrophages between the chicks infected with the wild-type strain and those infected with the *invA* mutant strain (p = 0.020 – S1 Table). This could be due to the fact that only two Δ*invA-*inoculated animals had infected monocytes-macrophages, while the percentages of monocytes-macrophages were similar for the five animals tested. Another interesting point is that the analyses of the infected areas (labeled + unlabeled) highlighted the high cell invasion rates of the gall bladder (Tab 1). Cumulating all experiments, after two days of infection, the median percentages of all infected cells (labeled and unlabeled) in the gall bladder were 2.23%, 1% and 3.65% depending on the strain inoculated, whereas in the spleen, for example, they were only 0.35%, 0.19% and 0.31%.

**Table 1.** Percentages of infected cells (labeled and unlabeled) according to the organ

|  | Spleen | Liver | Aortic vessels | Gall bladder |
| --- | --- | --- | --- | --- |
|  | Median (Q1; Q3) | Median (Q1; Q3) | Median (Q1; Q3) | Median (Q1; Q3) |
| WT | 0.35 (0.19; 0.53) | 0.07 (0.06; 0.10) | 0.64 (0.42; 2.49) | 2.23 (1.54; 2.82) |
| *ΔinvA* | 0.19 (0.14; 0.55) | 0.06 (0.04; 0.16) | 0.21 (0.12; 1.46) | 1.00 (0.52; 2.71) |
| 3Δ | 0.31 (0.26; 0.38) | 0.06 (0.04; 0.16) | 2.18 (1.18; 2.54) | 3.65 (2.39; 4.62) |

**Fig 6.**
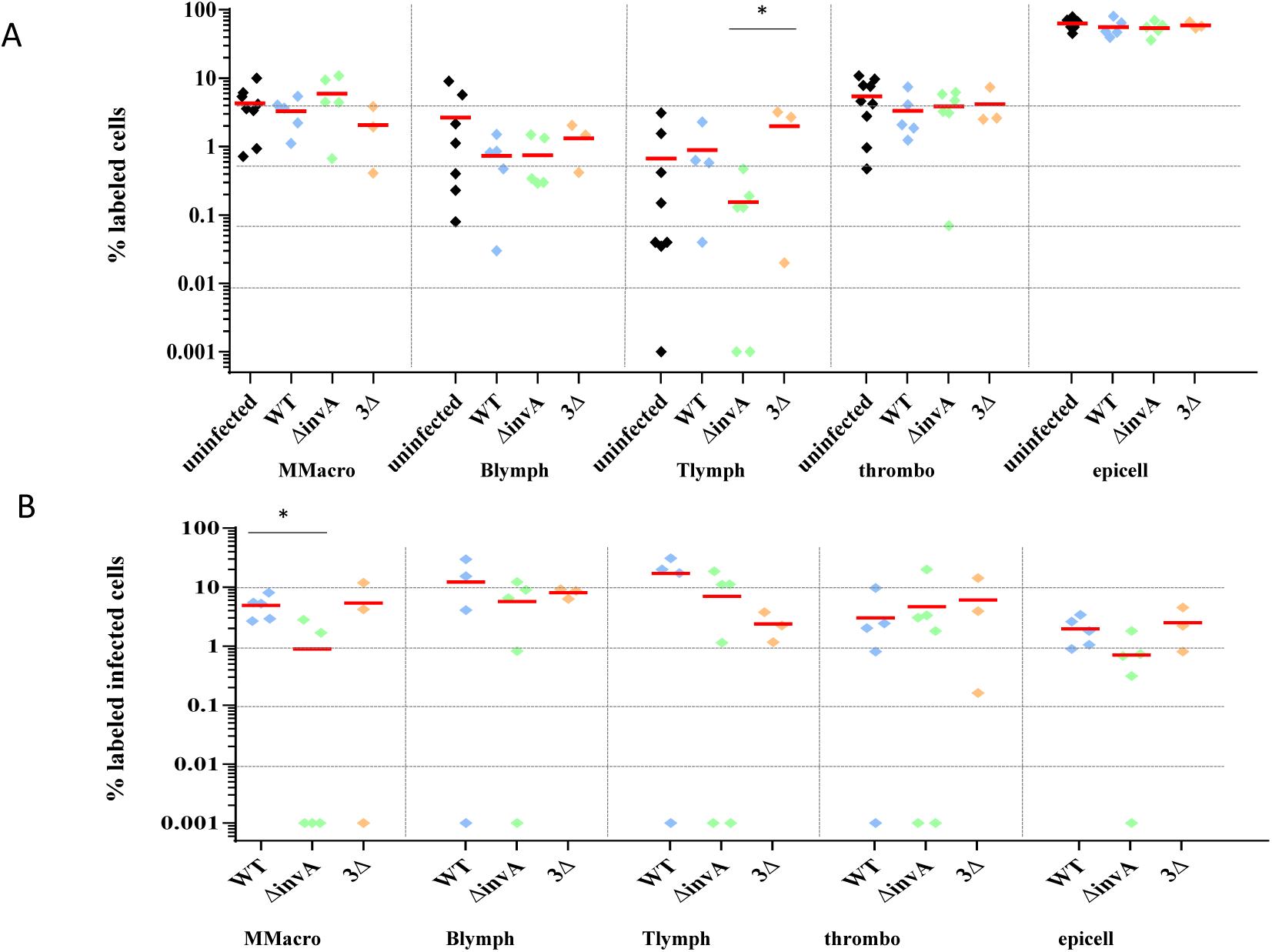
Percentage of identified and *Salmonella* infected cells in gall bladder. Five-day-old chicks were intraperitoneally inoculated with around 6.10^7^ CFU/chick with *S*. Typhimurium 14028 turboFP650 wild-type strain (WT 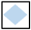), Δ*invA*::*kan* mutant strain (Δ*invA*; T3SS-1 defective 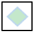) or the *ΔinvA::kan ΔpagN::cm Δrck* mutant strain (3Δ; T3SS-1, Rck, PagN defective 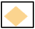). Two days post infection, animals were sacrificed and the different organs removed. Cells from uninfected animals of the same age were used as a control. After labeling with the corresponding antibodies, the percentages of macrophages-monocytes, B and T lymphocytes, thrombocytes and epithelial and endothelial cells were quantified by flow-cytometry. The percentage of labeled cells (A) and the percentage of labeled infected cells (B) are represented. All negative responses were scored at 0.001%. The medians are represented by a red dash. Asymptotic two-sample Fisher-Pitman permutation tests (One-Way-Test) were performed (R software). Significance was * p<0.05.

As this organ had never been described as a site of *Salmonella* colonization in chicks and as it is described as an organ that is important for *Salmonella* persistence in mice and humans (57, 58), we decided to observe the infected tissues using immunohistochemistry. Microscopic analysis shows that bacteria were located in the epithelium just above the *mucosa* of the gall bladder but also in the *lamina propria*, whatever the strain inoculated. Interestingly, when high numbers of *Salmonella* were detected, the epithelium was damaged while the structure of the gall bladder was well conserved when the tissue was only infected by a few bacteria (Fig 7).

**Fig 7.**
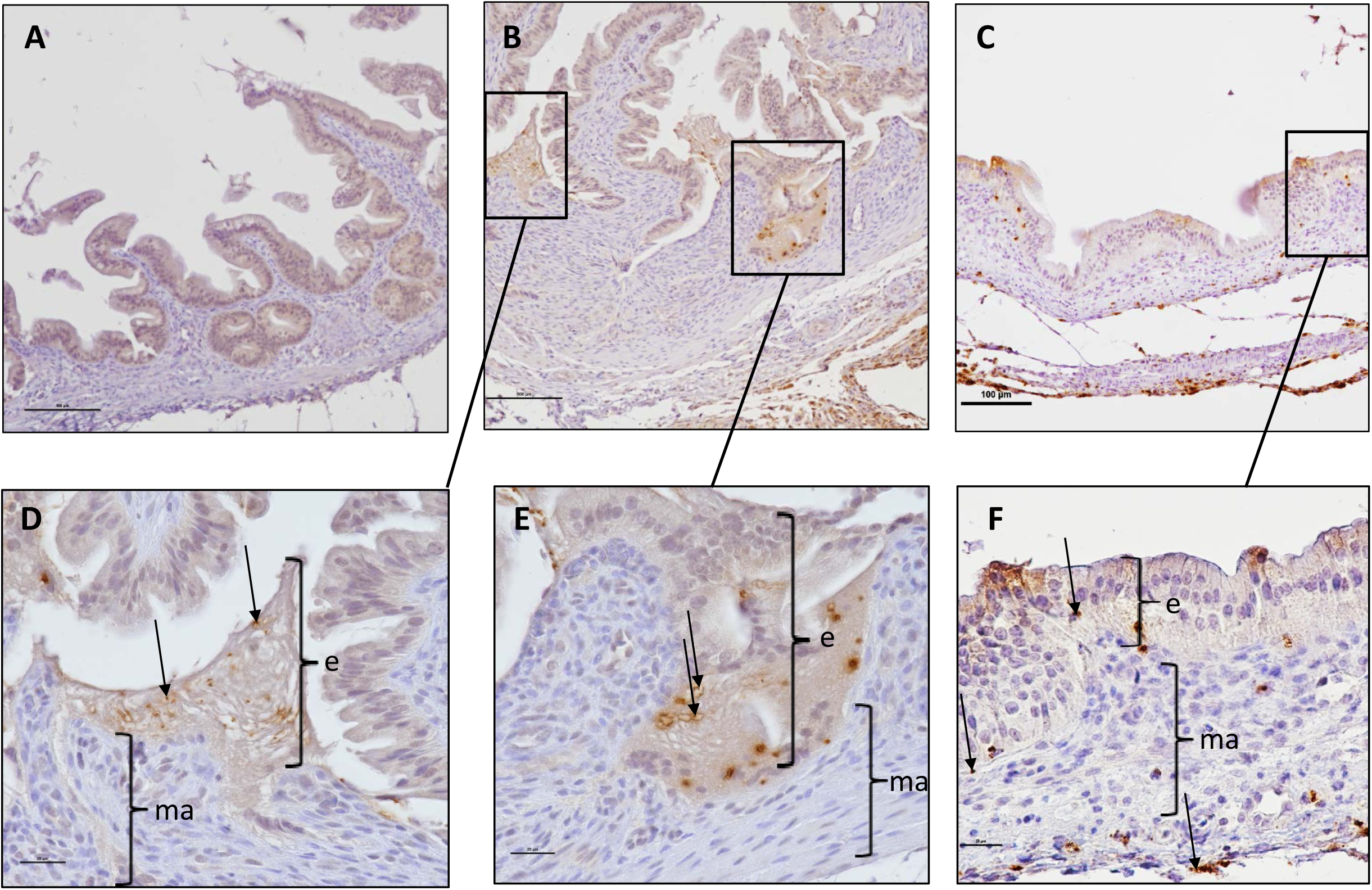
Immunohisto-chemistry of chick gall bladder infected with *S*. Typhimurium wild-type stain or with a mutant deleted of the three known invasion factors. Five-day-old chicks were intraperitoneally inoculated with around 6.10^7^ CFU/chick with *S*. Typhimurium 14028 turboFP650 wild-type strain (WT) or the *ΔinvA::kan ΔpagN::cm Δrck* mutant strain (3Δ). Two days post infection, animals were sacrificed. Gall bladders were removed and fixed in 4% buffered paraformaldehyde at 4°C for 24 h. Tissues were processed using routine methods, paraffin embedded, cut in sections (thickness, 5 µm), and stained with diaminobenzidine for IHC with HRP detection. The primary antibody was a rabbit anti-*Salmonella* O:4,5 lipopolysaccharide marker. Tissues were examined and photographed with a light microscope Eclipse 80i, Nikon with DXM 1200C digital camera (Nikon Instruments, Europe, Amsterdam, Netherlands) and NIS-Elements D Microscope Imaging Software. Tissues were counterstained in blue with Harris’ hematoxylin of and *Salmonella* were stained in brown with HRP detection. Representative pictures are presented. Bacteria are seen (↘) within the epithelium (e) and the mucosa (ma). Sections of a gall bladder of (A) an uninfected chick, (B, D and E) a chick infected by the wild-type strain, (C and F) a chick infected by the 3Δ mutant are represented.

The gall bladder is thus colonized by *Salmonella* after chicks are inoculated via an intraperitoneal route. These results show that oral inoculation of *Salmonella* is not necessary for gall bladder infection. The bacteria reached the gall bladder through the vasculature or the ducts that emanate from the liver. Menendez *et al*. obtained similar results in a mouse infection model (59). Indeed, they demonstrated that gallbladder colonization was not the result of *Salmonella* ascending directly from the gastrointestinal tract and their histological analyzes supported the idea that bacteria were discharged from the liver into the gall bladder *via* the bile. Concerning the infected cells in the chick gall bladder, monocytes-macrophages, B and T lymphocytes, thrombocytes and epithelial cells of this organ were all infected at a relatively high level compared to the other organs and epithelial cells were the most infected cell type.

Menendez *et. al* also observed in their mouse model that *Salmonella* localized preferentially within epithelial cells of the gallbladder. However, bacteria were rarely seen within the *lamina propria* (59). Our observations of some *S*. Typhimurium in the *mucosa* and *submucosa* and the identification of infected monocytes-macrophages, B and T lymphocytes, and thrombocytes demonstrate that epithelial cells are not the only cells infected in the gall bladder of chicks. Whether this result is restricted to chicks or not remains to be determined. Our results on our mutant strains are also different from that of Menendez *et al*. (59). Indeed, our mutants were shown to infect similar cells to the wild-type, while Menendez *et al*. did not observe their *inv*A mutant in the epithelial cells of the murine gall bladder, in contrast to their wild-type strain, suggesting that T3SS-1 could be required for *Salmonella* colonization of the gall bladder in mice but not in chicks.

The gall bladder is known to be an organ in which *S*. Typhi persist during chronic infections in humans, after forming a biofilm on gallstones (31, 58, 60). Models of chronic infection in mice have also been studied (57, 59, 61). In guinea pigs, although asymptomatic, *Salmonella* could be recovered in the gall bladder for up to 5 months post-infection (62). However, it is not known whether this organ could be relevant for the persistence of *Salmonella* in chicken.

### *Salmonella* Typhimurium is able to persist in the gall bladder independently of the T3SS1, Rck and PagN

As the previous results demonstrated that bacterial concentrations in the gall bladder were significant (Fig 1) and that *Salmonella* was able to infect several cell types in this organ, we verified whether this organ could be infected over the long term. To determine persistence in this organ, we infected chicks and monitored their colonization rate in the spleen and gall bladder for 36 days with slaughtering every 8 days. Bacteria were detected throughout the kinetics. No statistical differences of colonization could be observed in the spleen (Fig 8A) or in the gall bladder (Fig 8B), whatever the strain inoculated and the week of analysis. This work demonstrates for the first time, that *Salmonella* Typhimurium could invade the gall bladder of chicks at levels and durations similar to those observed in the spleen and thus, can be considered as a site of colonization in addition to the spleen and the liver in chicks. In mice, during chronic infection, mimicking human *S*. Typhi infection, the spleen and the gall bladder are considered as organs of persistence and gall bladder colonization presumably leads to re-infection of the intestine through bile secretion (46, 57, 63). Our results also demonstrated that, for the gall bladder to be infected, an oral route of infection is not necessary, as demonstrated by Menendez *et al*. in a mouse model (59). The role of this organ colonization needs to be analyzed further especially in relation to *Salmonella* intestinal colonization and persistence.

**Fig 8.**
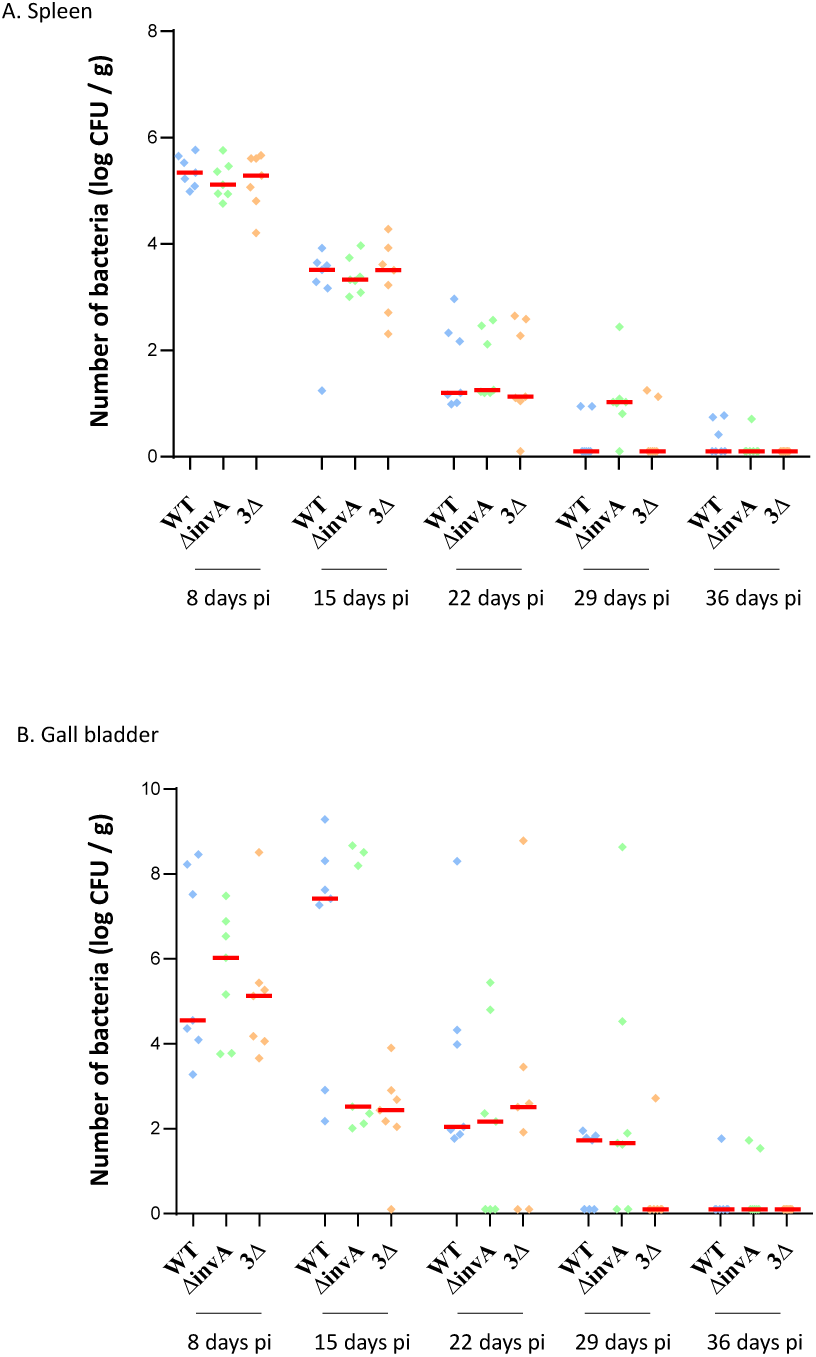
Persistence of *S*. Typhimurium in the spleen and in the gall bladder after intraperitoneal inoculation. Five-day-old chicks were intraperitoneally inoculated with around 3.10^7^ CFU/chick with *S*. Typhimurium 14028 turboFP650 wild-type (WT 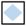), Δ*invA*::*kan* mutant strain (Δ*invA*; T3SS-1 defective 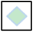) or the *ΔinvA::kan ΔpagN::cm Δrck* mutant strain (3Δ; T3SS-1, Rck, PagN defective 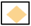). Each week, seven animals were sacrificed and their spleens and gall bladders removed. The kinetics of spleen (A) and gall bladder (B) colonization were followed each week for a period of 36 days. Results are expressed as number of bacteria (log CFU per g of organ). The medians are represented by a red dash. A Kruskal-Wallis test was conducted, followed by a Dunn’s multiple comparisons test (GraphPad Software). Significance was *p < 0.05.

## Conclusions

This work demonstrates for the first time, that *S*. Typhimurium can invade *in vivo* a large array of phagocytic and non-phagocytic cells of different organs and vessels in chicks. These cells are immune cells but also epithelial and endothelial cells as previously demonstrated *in vitro* with cell lines (30). Moreover, numerous unidentified cells were infected. This is due to the lack of antibodies in chicken to identify among others, dendritic cells, fibroblasts, and heterophils, which are also important cells for the spread of the bacterium (46, 64, 65). Development of new antibodies is required for further studies in chicks. Nevertheless, our results show a great difference between mice and chicks. In mice, phagocytic cells and especially macrophages are the main cells in which *Salmonella* replicate in the liver and spleen (41-43, 46). In chicks, the cell tropism in these organs, as well as in the gall bladder and vessels, is more diverse as *Salmonella* were found intracellularly in monocytes-macrophages but also in lymphocytes, endothelial cells and epithelial cells. Whether macrophages are the privileged localization of *Salmonella* or not in organs other than the liver and spleen in mice remains to be determined.

Even if *Salmonella* are able to invade numerous cells, specificity occurs depending on the organ. Indeed, for example, during a *Salmonella* infection, epithelial cells appeared more sensitive in the gall bladder than in the liver. In the same way, endothelial cells appeared more sensitive in the spleen than in the aortic vessels.

Surprisingly, the two mutant strains used in this study, i.e. a T3SS-1 mutant strain and a mutant strain defective for the three currently known invasion factors, were able to invade the same host cells as the wild-type strain. The fact that the triple mutant strain enters numerous host cells, *in vivo*, confirms our previous results suggesting the existence of unknown invasion factors. Indeed, we previously demonstrated that, despite the invalidation of the T3SS-1, Rck and PagN, *S*. Typhimurium remained able to invade, *in vitro*, some non-phagocytic cell lines of several animal and tissue origins at a similarly high level as the wild-type (30). However, we cannot conclude that the T3SS-1, PagN and Rck are not required for the invasion of chicken cells as a redundant role of the different invasion factors may occur. These two hypotheses are reinforced by our results of chicken infection demonstrating that chicks can be colonized at a higher level by the two mutant strains than by their wild-type parent after intraperitoneal inoculation. The absence of T3SS-1 requirement for chicken colonization has already been observed (18-20) and this fact can now be broadened to the PagN and Rck invasins. However, in order to demonstrate the redundant role or not of these entry factors, further studies are necessary in chicks and other animals. Altogether, these results are important in the understanding of the mechanisms of *Salmonella* pathogenesis, as it has been described that depending on the entry mechanism, both bacterial behavior and host response are different (28) and thus opening up new avenues of research. On the one hand, it raises the question as to whether the bacterial factors required for chicken cell invasion of systemic sites are still unknown and, if so, whether certain cell types are infected *in vivo* by a particular entry mechanism. On the other hand, could the known invasion factors in chicks be redundant? In addition, does this mean that because cells can be infected through multiple pathways in an organ, their response is multiple? Further studies involving Tnseq-mutant library screenings and single cell approaches would help to address these questions.

## Acknowledgements

We would like to thank Jérôme Trotereau (SPVB, ISP unit, INRAE Val de Loire) who participated in inoculating the chicks and the staff of the Experimental Platform for Infectious Diseases of Institut National de Recherche pour l’Agriculture, l’Alimentation et l’Environnement (PFIE, INRAE Val de Loire) for caring for the chicks and participating in the experiments, and also P. Quéré (3IMo, ISP unit, INRAE Val de Loire) for her advice on the labeling of the monocytes-macrophages. We are also grateful to T. Larcher (PAnTher APEX, INRAE – Oniris Nantes) for his advice and technical support.

## Funding

This work was supported by funding from the European Union’s Horizon 2020 Research and Innovation program under grant agreement No 773830: One Health European Joint Programme on Monitoring the gut Microbiota and Immune Response to Predict, Prevent, and Control zoonoses in humans and livestock in order to minimize the use of antimicrobials (MoMIR-PPC) https://onehealthejp.eu/jrp-momir/. A.R held an INRA/Region Centre doctoral fellowship.

## Figure legends

**Fig S1. Identification of labeled and infected labeled monocytes-macrophages using flow-cytometry**

The antibody allowing identification of monocytes-macrophages required a secondary Alexa Fluor™ 488 conjugated anti-mouse antibody. Flow cytometric analyses were performed with a BD LSR Fortessa™ X-20 (BD Biosciences, San Jose, CA, USA). BD FACSDiva™software (v 8.0.2) was used to analyze the cytometric data. For each sample, dot plots were analyzed. Debris was removed on the basis of morphological criteria, regions were defined on the basis of uninfected control samples and isotype-control staining. The intensity of green fluorescence (Alexa Fluor™ 488) was on the vertical axis, plotted against the intensity of red fluorescence (TurboFP650) on the horizontal axis. Labeled cells emitting a green fluorescence were detected in the upper part of the graph. Infected labeled cells emitting both types of fluorescence (green and red) were revealed by dots in the upper right-hand part of the graph. Unlabeled infected cells could also be seen in the lower right-hand part of the graph. Results were expressed as a percentage. Some examples are presented. Staining in the gall bladder is shown for the monocytes-macrophages.

**Fig S2. Identification of labeled and infected labeled thrombocytes using flow-cytometry**

The antibody allowing identification of thrombocytes required a secondary Alexa Fluor™ 488 conjugated anti-mouse antibody. Flow cytometric analyses were performed with a BD LSR Fortessa™ X-20 (BD Biosciences, San Jose, CA, USA). BD FACSDiva™ software (v 8.0.2) was used to analyze the cytometric data. For each sample, dot plots were analyzed. Debris was eliminated on the basis of morphological criteria, regions were set according to uninfected control samples and isotype-control staining. The intensity of green fluorescence (Alexa Fluor™ 488) is on the vertical axis, plotted against the intensity of red fluorescence (TurboFP650) on the horizontal axis. Labeled cells emitting a green fluorescence were detected in the upper part of the graph. Infected labeled cells emitting both types of fluorescence (green and red) were revealed by dots in the upper right-hand part of the graph. Unlabeled infected cells could also be seen in the lower right-hand part of the graph. Results were expressed as a percentage. Some examples are presented. Staining in the gall bladder is shown for the thrombocytes.

**Fig S3. Identification of labeled and infected labeled monocytes-macrophages using flow-cytometry**

The anti-Bu antibody, that allows B lymphocytes identification, was FITC conjugated. Flow cytometric analyses were performed with a BD LSR Fortessa™ X-20 (BD Biosciences, San Jose, CA, USA). BD FACSDiva™software (v 8.0.2) was used to analyze the cytometric data. For each sample, dot plots were analyzed. Debris was eliminated on the basis of morphological criteria, regions were set according to uninfected control samples and isotype-control staining. The intensity of green fluorescence (FITC) is on the vertical axis, plotted against the intensity of red fluorescence (TurboFP650) on the horizontal axis. Labeled cells emitting a green fluorescence were detected in the upper part of the graph. Infected labeled cells emitting both types of fluorescence (green and red) were revealed by dots in the upper right-hand part of the graph. Unlabeled infected cells could also be seen in the lower right-hand part of the graph. Results are expressed as percentages. Some examples are presented. Staining in the spleen is shown for the B lymphocytes.

**Fig S4. Identification of labeled and infected labeled T lymphocytes using flow-cytometry**

The anti-CD3 antibody, that allows T lymphocytes identification, was conjugated with FITC. Flow cytometric analyses were performed with a BD LSR Fortessa™ X-20 (BD Biosciences, San Jose, CA, USA). BD FACSDiva™ software (v 8.0.2) was used to analyze the cytometric data. For each sample, dot plots were analyzed. Debris was eliminated on the basis of morphological criteria, regions were set according to uninfected control samples and isotype-control staining. The intensity of green fluorescence (FITC) is on the vertical axis, plotted against the intensity of red fluorescence (TurboFP650) on the horizontal axis. Labeled cells emitting a green fluorescence were detected in the upper part of the graph. Infected labeled cells emitting both types of fluorescence (green and red) were revealed by dots in the upper right-hand part of the graph. Unlabeled infected cells could also be seen in the lower right-hand part of the graph. Results are expressed as percentages. Some examples are presented. Staining in the spleen is shown for the T lymphocytes.

**Fig S5. Identification of labeled and infected labeled epithelial cells using flow-cytometry**

The antibody allowing identification of epithelial cells required a secondary Alexa Fluor™ 488 conjugated anti-mouse antibody. Flow cytometric analyses were performed with a BD LSR Fortessa™ X-20 (BD Biosciences, San Jose, CA, USA). BD FACSDiva™ software (v 8.0.2) was used to analyze the cytometric data. For each sample, dot plots were analyzed. Debris was eliminated on the basis of morphological criteria, regions were set according to uninfected control samples and isotype-control staining. The intensity of green fluorescence (Alexa Fluor™ 488) is on the vertical axis, plotted against the intensity of red fluorescence (TurboFP650) on the horizontal axis. Labeled cells emitting a green fluorescence were detected in the upper part of the graph. Infected labeled cells emitting both types of fluorescence (green and red) were revealed by dots in the upper right-hand part of the graph. Unlabeled infected cells could also be seen in the lower right-hand part of the graph. Results are expressed as percentages. Some examples are presented. Staining in the liver is shown for epithelial cells.

**Fig S6. Identification of labeled and infected labeled endothelial cells using flow-cytometry**

The antibody allowing identification of endothelial cells required a secondary Alexa Fluor™ 488 conjugated anti-rabbit antibody. Flow cytometric analyses were performed with a BD LSR Fortessa™ X-20 (BD Biosciences, San Jose, CA, USA). BD FACSDiva™ software (v 8.0.2) was used to analyze the cytometric data. For each sample, dot plots were analyzed. Debris was eliminated on the basis of morphological criteria, regions were set according to uninfected control samples and isotype-control staining. The intensity of green fluorescence (Alexa Fluor™ 488) is on the vertical axis, plotted against the intensity of red fluorescence (TurboFP650) on the horizontal axis. Labeled cells emitting a green fluorescence were detected in the upper part of the graph. Infected labeled cells emitting both types of fluorescence (green and red) were revealed by dots in the upper right-hand part of the graph. Unlabeled infected cells could also be seen in the lower right-hand part of the graph. Results are expressed as percentages. Some examples are presented. Staining in the aortic vessels is shown for endothelial cells.

## Notes

### Competing Interest Statement

The authors have declared no competing interest.

## References

1. WHO. World Health Organization. Foodborne Disease Burden Epidemiology Reference Group. 2015. WHO estimates of the global burden of foodborne diseases. World Health Organization. 2017.

2. ECDC Ea. The European Union summary report on trends and sources of zoonoses, zoonotic agents and food-borne outbreaks in 2016. EFSA Journal. 2019;15(12):5077.

3. Ferrari RG, Rosario DKA, Cunha-Neto A, Mano SB, Figueiredo EES, Conte-Junior CA. Worldwide epidemiology of Salmonella serovars in animal-based foods: a meta-analysis. Appl Environ Microbiol. 2019;85(14): e00591–19.

4. Rabinowitz PM, Conti LA. Human-clinical-medicine: clinical approaches to zoonoses, toxicants and other shared health risks. 1st ed. Saunders Maryland Heights, MD, USA;2009.

5. Knight-Jones TJ, Mylrea GE, Kahn S. Animal production food safety: priority pathogens for standard setting by the World Organisation for Animal Health. Rev Sci Tech. 2010;29(3):523–35.

6. Menanteau P, Kempf F, Trotereau J, Virlogeux-Payant I, Gitton E, Dalifard J, et al. Role of systemic infection, cross contaminations and super-shedders in Salmonella carrier state in chicken. Environ Microbiol. 2018;20(9):3246–60.

7. Ly KT, Casanova JE. Mechanisms of Salmonella entry into host cells. Cell Microbiol. 2007;9(9):2103–11.

8. Heffernan EJ, Wu L, Louie J, Okamoto S, Fierer J, Guiney DG. Specificity of the complement resistance and cell association phenotypes encoded by the outer membrane protein genes rck from Salmonella Typhimurium and ail from Yersinia enterocolitica. Infect Immun. 1994;62(11):5183–6.

9. Rosselin M, Virlogeux-Payant I, Roy C, Bottreau E, Sizaret PY, Mijouin L, et al. Rck of Salmonella enterica, subspecies enterica serovar Enteritidis, mediates zipper-like internalization. Cell Res. 2010;20(6):647–64.

10. Lambert MA, Smith SG. The PagN protein of Salmonella enterica serovar Typhimurium is an adhesin and invasin. BMC Microbiol. 2008;8:142.

11. Lambert MA, Smith SG. The PagN protein mediates invasion via interaction with proteoglycan. FEMS Microbiol Lett. 2009;297(2):209–16.

12. Wiedemann A, Mijouin L, Ayoub MA, Barilleau E, Canepa S, Teixeira-Gomes AP, et al. Identification of the epidermal growth factor receptor as the receptor for Salmonella Rck-dependent invasion. FASEB journal : official publication of the Federation of American Societies for Experimental Biology. 2016;30(12):4180–91.

13. Wallis TS, Galyov EE. Molecular basis of Salmonella-induced enteritis. Mol Microbiol. 2000;36(5):997–1005.

14. Jneid B, Moreau K, Plaisance M, Rouaix A, Dano J, Simon S. Role of T3SS-1 SipD Protein in Protecting Mice against Non-typhoidal Salmonella Typhimurium. PLoS Negl Trop Dis. 2016;10(12):e0005207.

15. Galan JE, Curtiss R, 3rd. Cloning and molecular characterization of genes whose products allow Salmonella Typhimurium to penetrate tissue culture cells. PNAS. 1989;86(16):6383–7.

16. Murray RA, Lee CA. Invasion genes are not required for Salmonella enterica serovar Typhimurium to breach the intestinal epithelium: evidence that Salmonella pathogenicity island 1 has alternative functions during infection. Infect Immun. 2000;68(9):5050–5.

17. Hapfelmeier S, Stecher B, Barthel M, Kremer M, Muller AJ, Heikenwalder M, et al. The Salmonella pathogenicity island (SPI)-2 and SPI-1 type III secretion systems allow Salmonella serovar Typhimurium to trigger colitis via MyD88-dependent and MyD88-independent mechanisms. J Immunol. 2005;174(3):1675–85.

18. Sivula CP, Bogomolnaya LM, Andrews-Polymenis HL. A comparison of cecal colonization of Salmonella enterica serotype Typhimurium in white leghorn chicks and Salmonella-resistant mice. BMC Microbiol. 2008;8:182.

19. Rychlik I, Karasova D, Sebkova A, Volf J, Sisak F, Havlickova H, et al. Virulence potential of five major pathogenicity islands (SPI-1 to SPI-5) of Salmonella enterica serovar Enteritidis for chickens. BMC Microbiol. 2009;9:268.

20. Desin TS, Lam PK, Koch B, Mickael C, Berberov E, Wisner AL, et al. Salmonella enterica serovar Enteritidis pathogenicity island 1 is not essential for but facilitates rapid systemic spread in chickens. Infect Immun. 2009;77(7):2866–75.

21. Hu Q, Coburn B, Deng W, Li Y, Shi X, Lan Q, et al. Salmonella enterica serovar Senftenberg human clinical isolates lacking SPI-1. J Clin Microbiol. 2008;46(4):1330–6.

22. Suez J, Porwollik S, Dagan A, Marzel A, Schorr YI, Desai PT, et al. Virulence gene profiling and pathogenicity characterization of non-typhoidal Salmonella accounted for invasive disease in humans. PLoS One. 2013;8(3):e58449.

23. Conner CP, Heithoff DM, Julio SM, Sinsheimer RL, Mahan MJ. Differential patterns of acquired virulence genes distinguish Salmonella strains. PNAS. 1998;95(8):4641–5.

24. Dyszel JL, Smith JN, Lucas DE, Soares JA, Swearingen MC, Vross MA, et al. Salmonella enterica serovar Typhimurium can detect acyl homoserine lactone production by Yersinia enterocolitica in mice. J Bacteriol. 2010;192(1):29–37.

25. Ghosh S, Chakraborty K, Nagaraja T, Basak S, Koley H, Dutta S, et al. An adhesion protein of Salmonella enterica serovar Typhi is required for pathogenesis and potential target for vaccine development. PNAS. 2011;108(8):3348–53.

26. Tundup S, Kandasamy M, Perez JT, Mena N, Steel J, Nagy T, et al. Endothelial cell tropism is a determinant of H5N1 pathogenesis in mammalian species. PLoS Pathog. 2017;13(3):e1006270.

27. Pereira SS, Trindade S, De Niz M, Figueiredo LM. Tissue tropism in parasitic diseases. Open Biol. 2019;9:190036.

28. Miao EA, Mao DP, Yudkovsky N, Bonneau R, Lorang CG, Warren SE, et al. Innate immune detection of the type III secretion apparatus through the NLRC4 inflammasome. PNAS. 2010;107(7):3076–80.

29. Ryan KJ, Ray CGE. Sherris Medical Microbiology: an introduction to infectious disease (fourth edition). New York: McGraw-Hill, USA; 2004.

30. Roche SM, Holbert S, Trotereau J, Schaeffer S, Georgeault S, Virlogeux-Payant I, et al. Salmonella Typhimurium invalidated for the three currently known invasion factors keeps its ability to invade several cell models. Front Cell Infect Microbiol. 2018;8:273.

31. Gonzalez-Escobedo G, Marshall JM, Gunn JS. Chronic and acute infection of the gall bladder by Salmonella Typhi: understanding the carrier state. Nat Rev Microbiol. 2011;9(1):9–14.

32. Galan JE. Salmonella interactions with host cells: type III secretion at work. Annu Rev Cell Dev Biol. 2001;17:53–86.

33. Coburn B, Sekirov I, Finlay BB. Type III secretion systems and disease. Clin Microbiol Rev. 2007;20(4):535–49.

34. McGhie EJ, Brawn LC, Hume PJ, Humphreys D, Koronakis V. Salmonella takes control: effector-driven manipulation of the host. Curr Opin Microbiol. 2009;12(1):117–24.

35. Jones MA, Hulme SD, Barrow PA, Wigley P. The Salmonella pathogenicity island 1 and Salmonella pathogenicity island 2 type III secretion systems play a major role in pathogenesis of systemic disease and gastrointestinal tract colonization of Salmonella enterica serovar Typhimurium in the chicken. Avian Pathol. 2007;36(3):199–203.

36. Pavlova B, Volf J, Ondrackova P, Matiasovic J, Stepanova H, Crhanova M, et al. SPI-1-encoded type III secretion system of Salmonella enterica is required for the suppression of porcine alveolar macrophage cytokine expression. Vet Res. 2011;42:16.

37. Sekirov I, Gill N, Jogova M, Tam N, Robertson M, de Llanos R, et al. Salmonella SPI-1-mediated neutrophil recruitment during enteric colitis is associated with reduction and alteration in intestinal microbiota. Gut microbes. 2010;1(1):30–41.

38. Zhao X, Tang X, Guo N, An Y, Chen X, Shi C, et al. Biochanin a enhances the defense against Salmonella enterica infection through AMPK/ULK1/mTOR-mediated autophagy and extracellular traps and reversing SPI-1-dependent macrophage (MPhi) M2 polarization. Front Cell Infect Microbiol. 2018;8:318.

39. Lou L, Zhang P, Piao R, Wang Y. Salmonella Pathogenicity Island 1 (SPI-1) and its complex regulatory network. Front Cell Infect Microbiol. 2019;9:270.

40. Elsheimer-Matulova M, Varmuzova K, Kyrova K, Havlickova H, Sisak F, Rahman M, et al. phoP, SPI1, SPI2 and aroA mutants of Salmonella Enteritidis induce a different immune response in chickens. Vet Res. 2015;46:96.

41. Richter-Dahlfors A, Buchan AM, Finlay BB. Murine salmonellosis studied by confocal microscopy: Salmonella Typhimurium resides intracellularly inside macrophages and exerts a cytotoxic effect on phagocytes in vivo. J Exp Med. 1997;186(4):569–80.

42. Salcedo SP, Noursadeghi M, Cohen J, Holden DW. Intracellular replication of Salmonella Typhimurium strains in specific subsets of splenic macrophages in vivo. Cell Microbiol. 2001;3(9):587–97.

43. Sheppard M, Webb C, Heath F, Mallows V, Emilianus R, Maskell D, et al. Dynamics of bacterial growth and distribution within the liver during Salmonella infection. Cell Microbiol. 2003;5(9):593–600.

44. Geddes K, Cruz F, Heffron F. Analysis of cells targeted by Salmonella type III secretion in vivo. PLoS Pathog. 2007;3(12):e196.

45. Thone F, Schwanhausser B, Becker D, Ballmaier M, Bumann D. FACS-isolation of Salmonella-infected cells with defined bacterial load from mouse spleen. J Microbiol Methods. 2007;71(3):220–4.

46. Watson KG, Holden DW. Dynamics of growth and dissemination of Salmonella in vivo. Cell Microbiol. 2010;12(10):1389–97.

47. Mastroeni P, Grant A, Restif O, Maskell D. A dynamic view of the spread and intracellular distribution of Salmonella enterica. Nat Rev Microbiol. 2009;7(1):73–80.

48. Jeurissen SH. Structure and function of the chicken spleen. Res Immunol. 1991;142(4):352–5.

49. Fields PI, Swanson RV, Haidaris CG, Heffron F. Mutants of Salmonella Typhimurium that cannot survive within the macrophage are avirulent. PNAS. 1986;83(14):5189–93.

50. Zaefarian F, Abdollahi MR, Cowieson A, Ravindran V. Avian Liver: The Forgotten Organ. Animals (Basel). 2019;9(2).

51. Hu T, Wu Z, Bush SJ, Freem L, Vervelde L, Summers KM, et al. Characterization of subpopulations of chicken mononuclear phagocytes that express TIM4 and CSF1R. J Immunol. 2019;202(4):1186–99.

52. Mehal WZ, Azzaroli F, Crispe IN. Immunology of the healthy liver: old questions and new insights. Gastroenterology. 2001;120(1):250–60.

53. Wang Y, Zhang C. The roles of liver-resident lymphocytes in liver diseases. Front Immunol. 2019;10:1582.

54. Erdogan S. The branching of the aortic arch in the Eurasian bittern (Botaurus stellaris, Linnaeus 1758). Vet Med. 2012;57(5):239–44.

55. Tucker WD, Arora Y, Mahajan K. Anatomy, Blood Vessels. In: StatPearls (Internet). Treasure Island (FL): StatPearls Publishing 2020, PMID: 2922226.

56. Iqbal J, Bhutto AL, Shah MG, Lochi GM, Hayat S, Ali N, et al. Gross anatomical and histological studies on the liver of broiler. J Appl Environ Biol Sci. 2014;4(12):284–95.

57. Crawford RW, Rosales-Reyes R, Ramirez-Aguilar Mde L, Chapa-Azuela O, Alpuche-Aranda C, Gunn JS. Gallstones play a significant role in Salmonella spp. gallbladder colonization and carriage. PNAS. 2010;107(9):4353–8.

58. Di Domenico EG, Cavallo I, Pontone M, Toma L, Ensoli F. Biofilm producing Salmonella Typhi: chronic colonization and development of gallbladder cancer. Int J Mol Sci. 2017;18(9): 1887.

59. Menendez A, Arena ET, Guttman JA, Thorson L, Vallance BA, Vogl W, et al. Salmonella infection of gallbladder epithelial cells drives local inflammation and injury in a model of acute typhoid fever. J Infect Dis. 2009;200(11):1703–13.

60. Gunn JS, Marshall JM, Baker S, Dongol S, Charles RC, Ryan ET. Salmonella chronic carriage: epidemiology, diagnosis, and gallbladder persistence. Trends Microbiol. 2014;22(11):648–55.

61. Scanu T, Spaapen RM, Bakker JM, Pratap CB, Wu LE, Hofland I, et al. Salmonella manipulation of host signaling pathways provokes cellular transformation associated with gallbladder carcinoma. Cell Host Microbe. 2015;17(6):763–74.

62. Lavergne GM, James HF, Martineau C, Diena BB, Lior H. The guinea pig as a model for the asymptomatic human typhoid carrier. Lab Anim Sci. 1977;27(5 Pt 2):806–16.

63. Gonzalez-Escobedo G, Gunn JS. Gallbladder epithelium as a niche for chronic Salmonella carriage. Infect Immun. 2013;81(8):2920–30.

64. van Dijk A, Tersteeg-Zijderveld MH, Tjeerdsma-van Bokhoven JL, Jansman AJ, Veldhuizen EJ, Haagsman HP. Chicken heterophils are recruited to the site of Salmonella infection and release antibacterial mature Cathelicidin-2 upon stimulation with LPS. Mol Immunol. 2009;46(7):1517–26.

65. Aiastui A, Pucciarelli MG, Garcia-del Portillo F. Salmonella enterica serovar Typhimurium invades fibroblasts by multiple routes differing from the entry into epithelial cells. Infect Immun. 2010;78(6):2700–13.

66. Valdivia RH, Falkow S. Bacterial genetics by flow cytometry: rapid isolation of Salmonella Typhimurium acid-inducible promoters by differential fluorescence induction. Mol Microbiol. 1996;22(2):367–78.

67. Casadaban MJ, Cohen SN. Analysis of gene control signals by DNA fusion and cloning in Escherichia coli. J Mol Biol. 1980;138(2):179–207.

68. Mast J, Goddeeris BM, Peeters K, Vandesande F, Berghman LR. Characterisation of chicken monocytes, macrophages and interdigitating cells by the monoclonal antibody KUL01. Vet Immunol Immunopathol. 1998;61(2-4):343–57.

69. Chen CL, Ager LL, Gartland GL, Cooper MD. Identification of a T3/T cell receptor complex in chickens. J Exp Med. 1986;164(1):375–80.

70. Rothwell CJ, Vervelde, L., Davison, T.F. Identification of Bu-1 alloantigens using the monoclonal antibody AV20. 1996. In: Poultry Immunology [Internet]. Carfax, Abindon, UK.

71. Lacoste-Eleaume AS, Bleux C, Quere P, Coudert F, Corbel C, Kanellopoulos-Langevin C. Biochemical and functional characterization of an avian homolog of the integrin GPIIb-IIIa present on chicken thrombocytes. Exp Cell Res. 1994;213(1):198–209.

72. Gallin WJ, Edelman GM, Cunningham BA. Characterization of L-CAM, a major cell adhesion molecule from embryonic liver cells. PNAS. 1983;80(4):1038–42.

