## Supplemental Figure 1 for "A large panel of chicken cells are invaded *in vivo* by *Salmonella* Typhimurium even when depleted of all known invasion factors"

### Slide 1
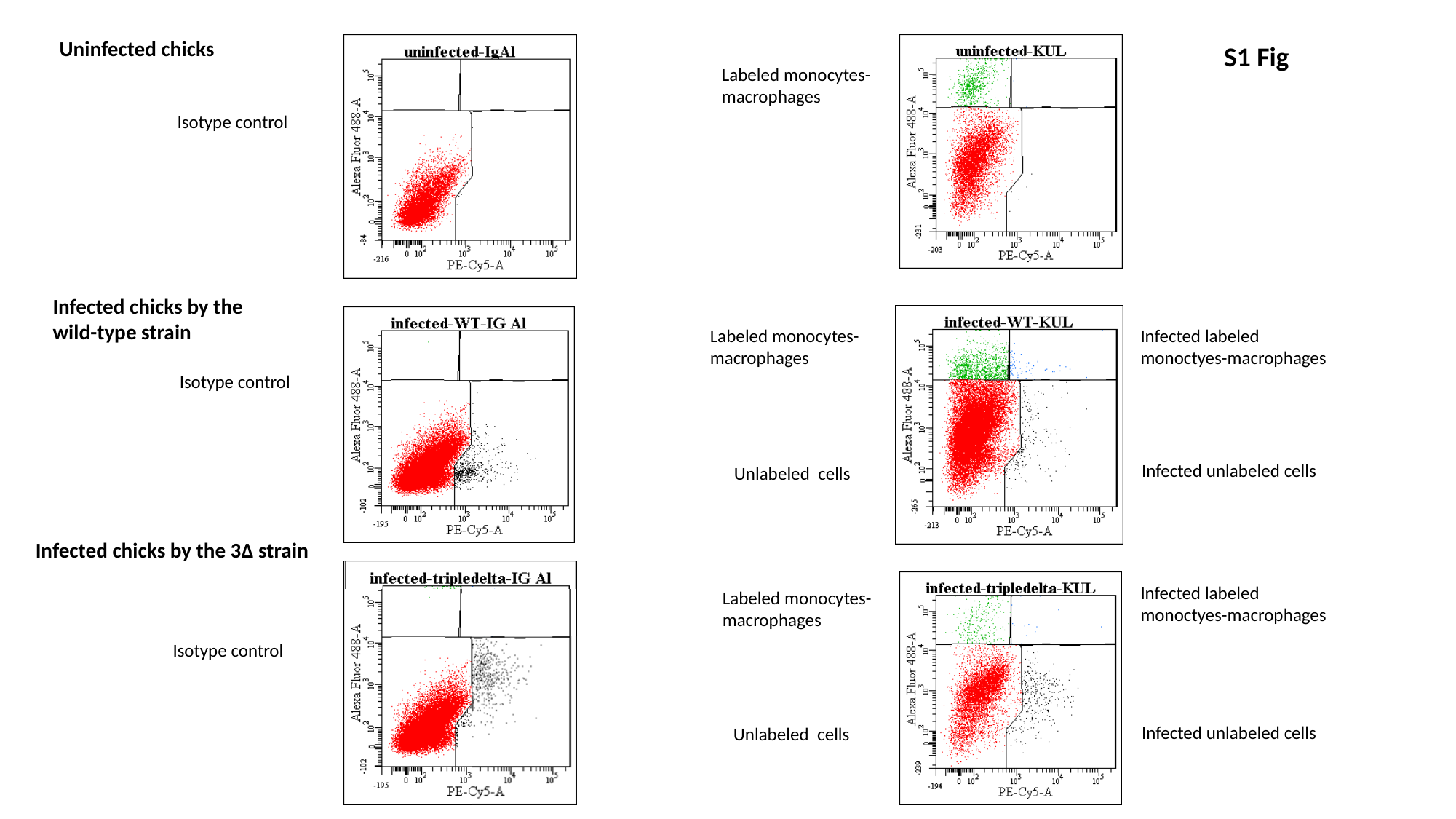

Uninfected chicks
Labeled monocytes-macrophages
Isotype control
Infected chicks by the wild-type strain
Infected labeled monoctyes-macrophages
Isotype control
Infected unlabeled cells
Unlabeled cells
Infected chicks by the 3Δ strain
Isotype control
Infected unlabeled cells
Unlabeled cells
S1 Fig
Labeled monocytes-macrophages
Infected labeled monoctyes-macrophages
Labeled monocytes-macrophages
