## Supplemental Figure 2 for "A large panel of chicken cells are invaded *in vivo* by *Salmonella* Typhimurium even when depleted of all known invasion factors"

### Slide 1
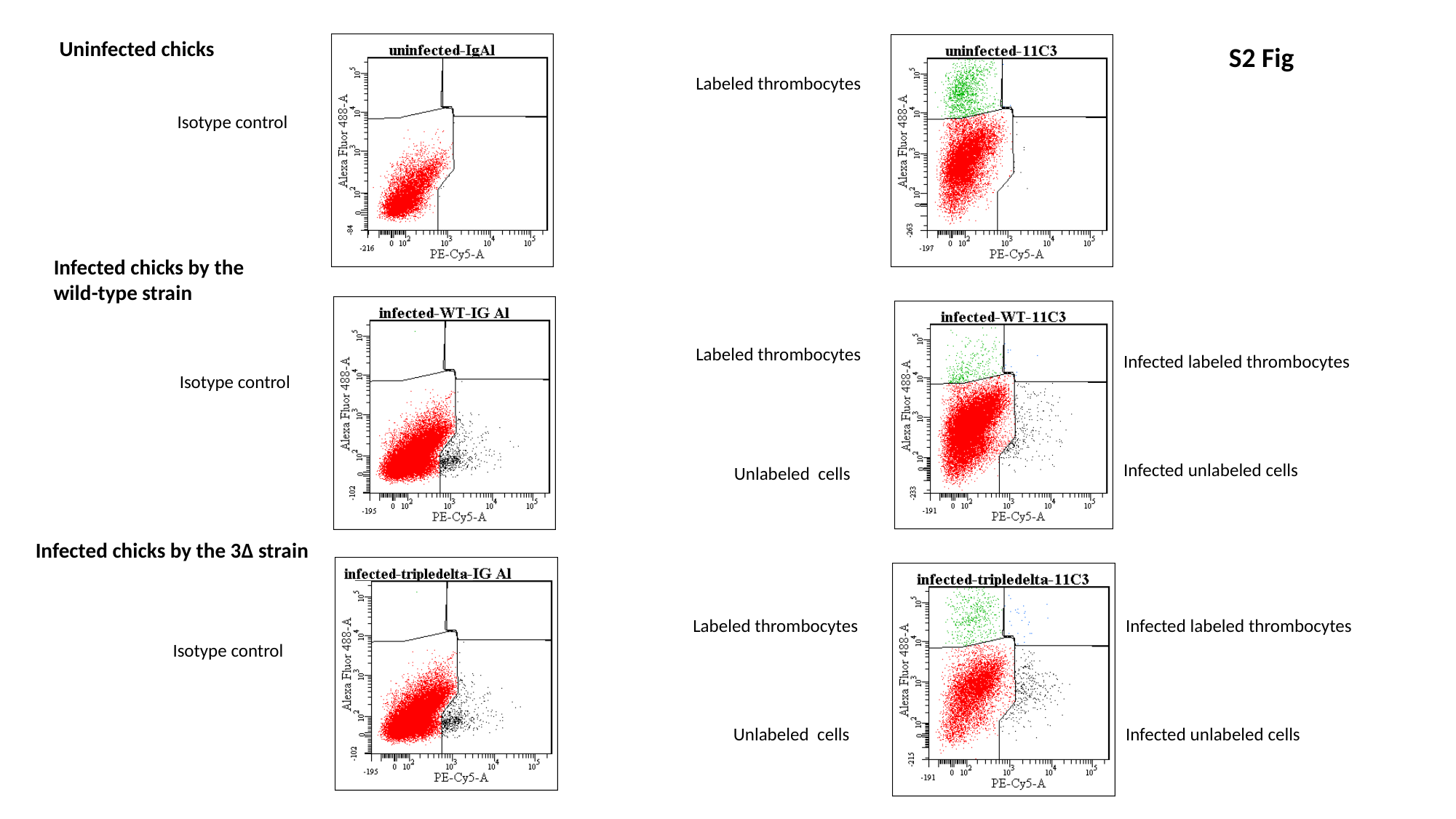

Uninfected chicks
Labeled thrombocytes
Isotype control
Infected chicks by the wild-type strain
Labeled thrombocytes
Infected labeled thrombocytes
Isotype control
Infected unlabeled cells
Unlabeled cells
Infected chicks by the 3Δ strain
Labeled thrombocytes
Infected labeled thrombocytes
Isotype control
Unlabeled cells
Infected unlabeled cells
S2 Fig
