## Supplemental Figure 3 for "A large panel of chicken cells are invaded *in vivo* by *Salmonella* Typhimurium even when depleted of all known invasion factors"

### Slide 1
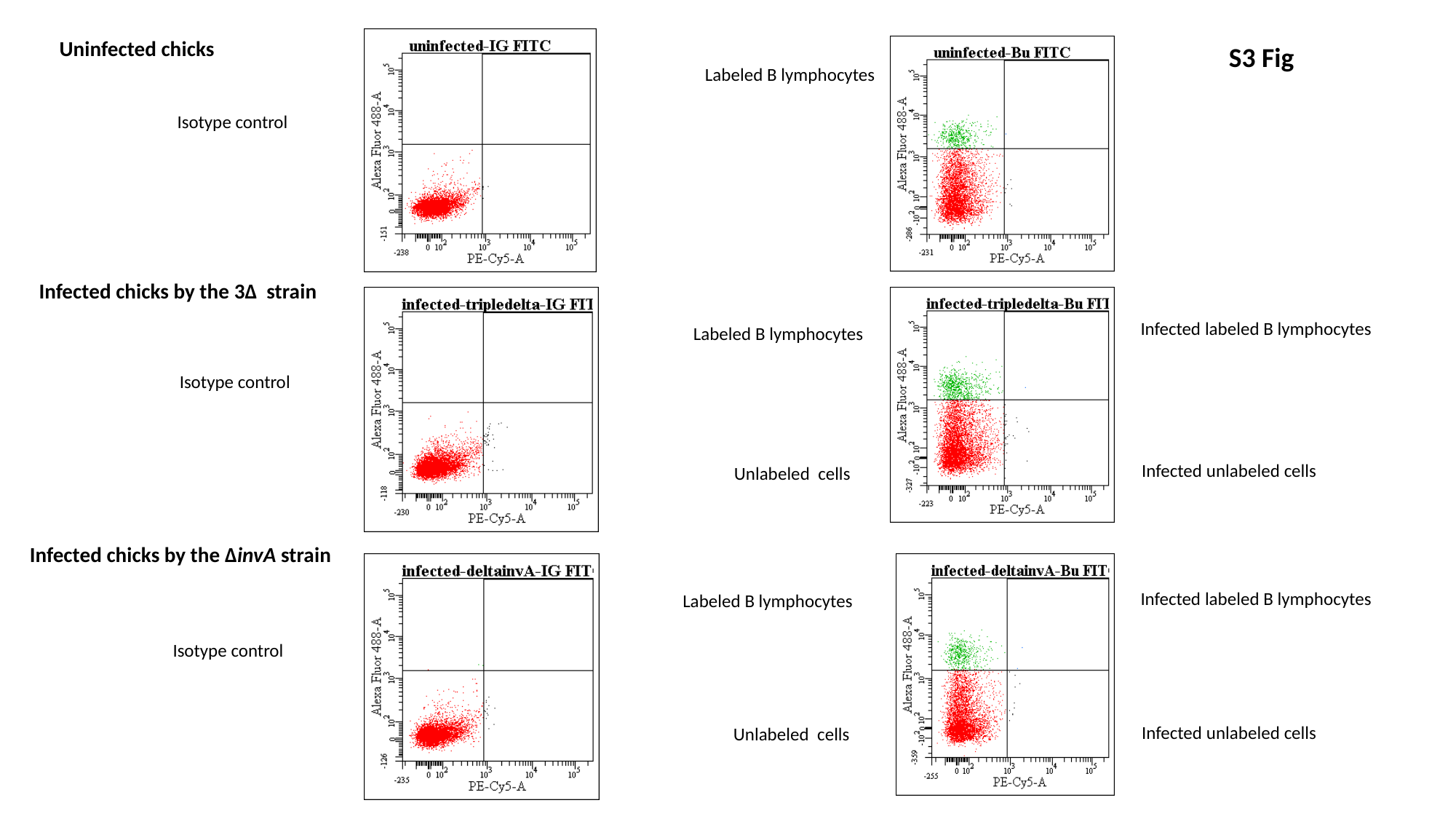

Uninfected chicks
Labeled B lymphocytes
Isotype control
Infected chicks by the 3Δ strain
Infected labeled B lymphocytes
Isotype control
Infected unlabeled cells
Unlabeled cells
Infected chicks by the ΔinvA strain
Isotype control
Infected unlabeled cells
Unlabeled cells
S3 Fig
Labeled B lymphocytes
Infected labeled B lymphocytes
Labeled B lymphocytes
