## Supplemental Figure 4 for "A large panel of chicken cells are invaded *in vivo* by *Salmonella* Typhimurium even when depleted of all known invasion factors"

### Slide 1
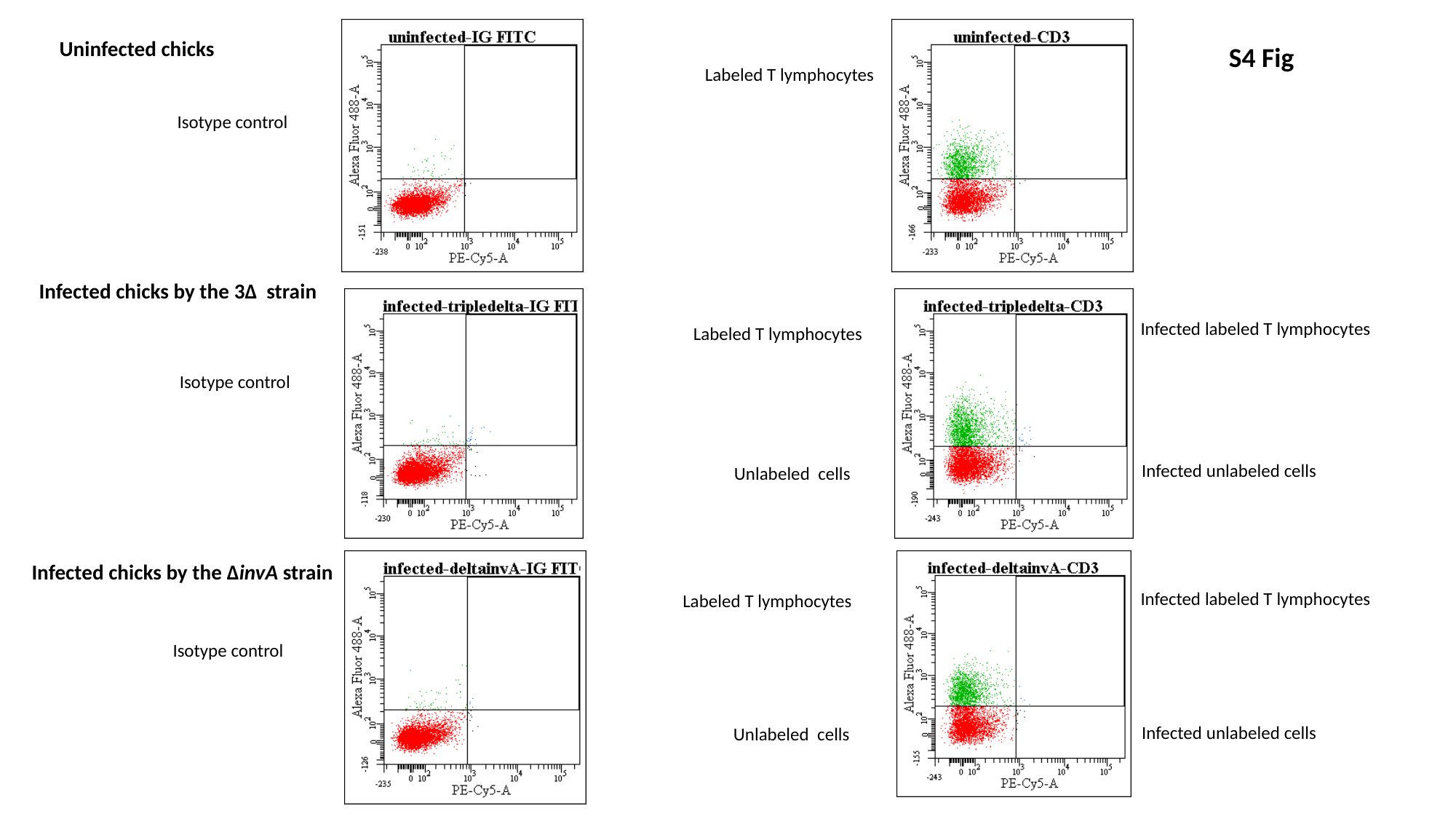

Uninfected chicks
Labeled T lymphocytes
Isotype control
Infected chicks by the 3Δ strain
Infected labeled T lymphocytes
Isotype control
Infected unlabeled cells
Unlabeled cells
Infected chicks by the ΔinvA strain
Isotype control
Infected unlabeled cells
Unlabeled cells
S4 Fig
Labeled T lymphocytes
Infected labeled T lymphocytes
Labeled T lymphocytes
