## Supplemental Figure 5 for "A large panel of chicken cells are invaded *in vivo* by *Salmonella* Typhimurium even when depleted of all known invasion factors"

### Slide 1
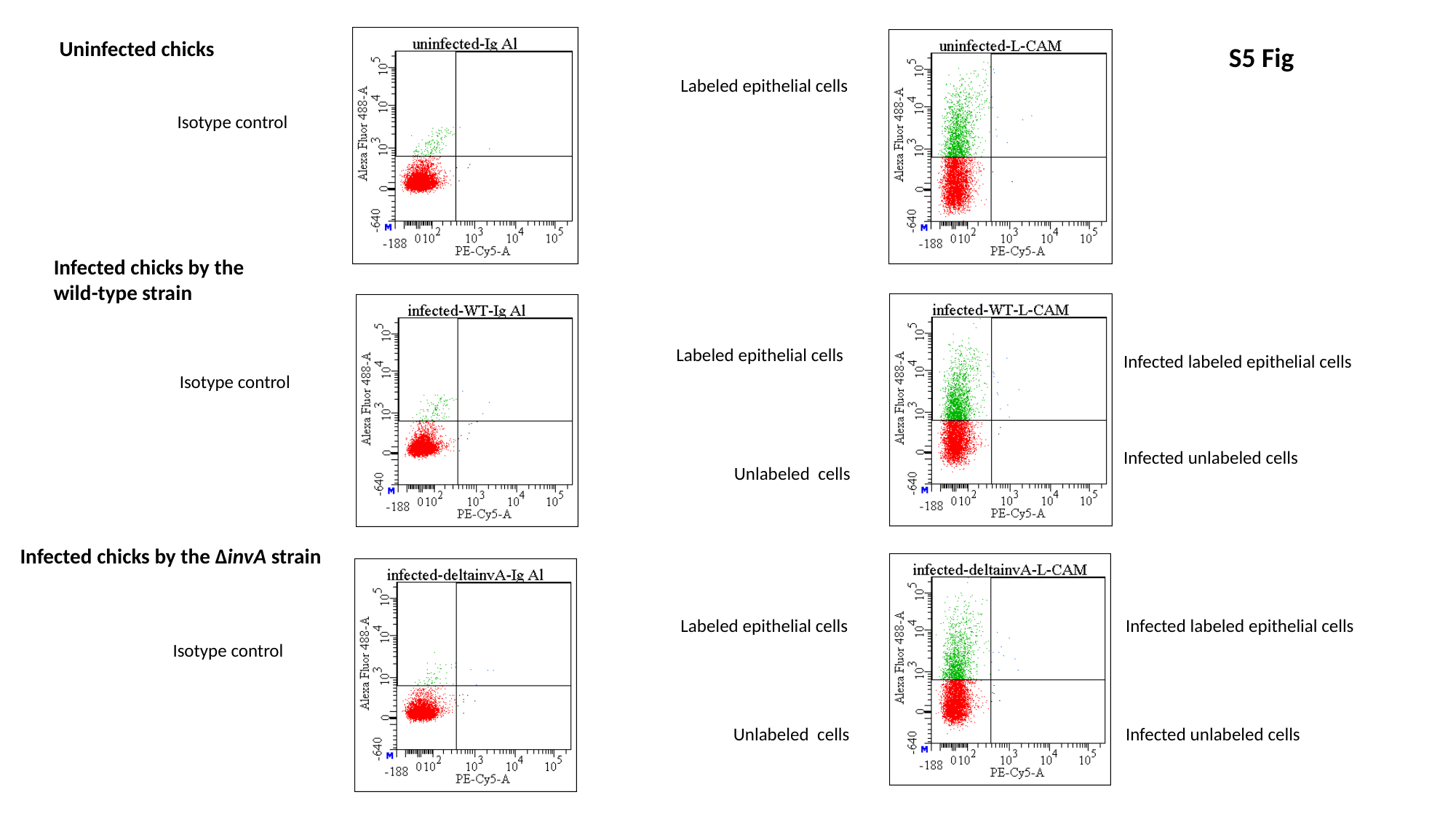

Uninfected chicks
Labeled epithelial cells
Isotype control
Infected chicks by the wild-type strain
Labeled epithelial cells
Infected labeled epithelial cells
Isotype control
Infected unlabeled cells
Unlabeled cells
Infected chicks by the ΔinvA strain
Labeled epithelial cells
Infected labeled epithelial cells
Isotype control
Unlabeled cells
Infected unlabeled cells
S5 Fig
