## Supplemental Figure 6 for "A large panel of chicken cells are invaded *in vivo* by *Salmonella* Typhimurium even when depleted of all known invasion factors"

### Slide 1
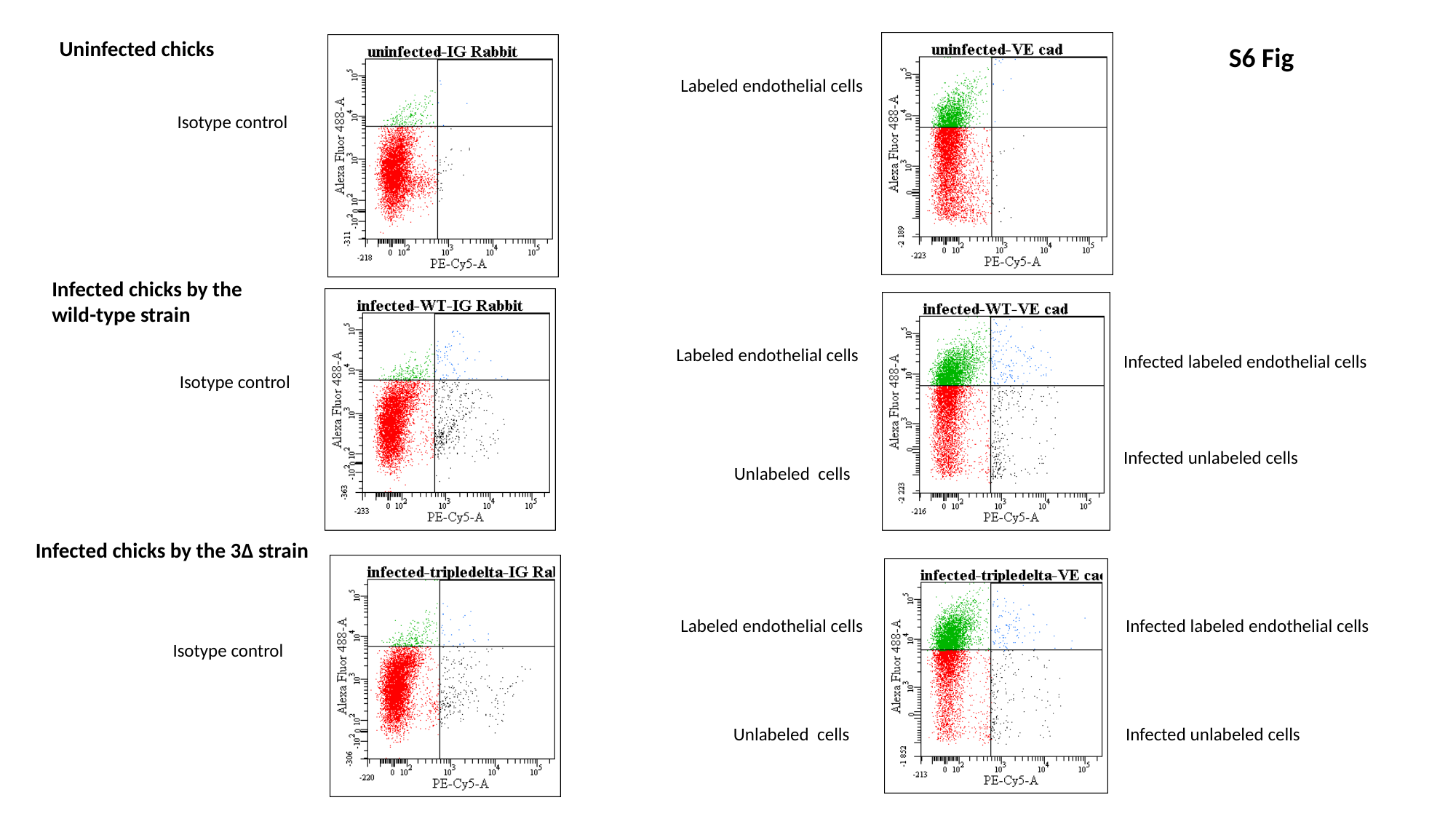

Uninfected chicks
Labeled endothelial cells
Isotype control
Infected chicks by the wild-type strain
Labeled endothelial cells
Infected labeled endothelial cells
Isotype control
Infected unlabeled cells
Unlabeled cells
Infected chicks by the 3Δ strain
Labeled endothelial cells
Infected labeled endothelial cells
Isotype control
Unlabeled cells
Infected unlabeled cells
S6 Fig
