## Supplemental Table 1 for "A large panel of chicken cells are invaded *in vivo* by *Salmonella* Typhimurium even when depleted of all known invasion factors"

S1 Table. Statistical analyzes of the cytometric results

|  | Cells |  | MMacro |  | Blymph | T lymph |  | thrombo |  | epicell |  | endocell |
| --- | --- | --- | --- | --- | --- | --- | --- | --- | --- | --- | --- | --- |
| Spleen | Labeled cells | uninfected-WT | 0.623 |  | 0.398 | 0.513 |  | **0.049** | ***** |  |  | 0.541 |
|  |  | uninfected-Δ*invA* | 0.222 |  | 0.750 | 0.972 |  | 0.053 |  |  |  | 0.591 |
|  |  | uninfected-3Δ | 0.207 |  | 0.154 | 0.150 |  | 0.069 |  |  |  | 0.437 |
|  |  | WT-Δ*invA* | 0.318 |  | 0.350 | 0.527 |  | 0.844 |  |  |  | 0.197 |
|  |  | WT-3Δ | 0.285 |  | 0.461 | 0.388 |  | 0.873 |  |  |  | 0.752 |
|  |  | Δ*invA*-3Δ | 0.647 |  | 0.160 | 0.186 |  | 0.186 |  |  |  | 0.143 |
|  | Infected labeled cells | WT-Δ*invA* | 0.943 |  | 0.969 | 0.244 |  | 0.605 |  |  |  | 0.954 |
|  |  | WT-3Δ | 0.534 |  | 0.264 | 0.232 |  | 0.687 |  |  |  | 0.539 |
|  |  | Δ*invA*-3Δ | 0.658 |  | 0.478 | 0.576 |  | 0.913 |  |  |  | 0.686 |
| Liver | Labeled cells | uninfected-WT | 0.936 |  | 0.371 | 0.504 |  | 0.101 |  | 0.854 |  | 0.350 |
|  |  | uninfected-Δ*invA* | 0.900 |  | 0.928 | 0.537 |  | 0.537 |  | **0.039** | * | 0.722 |
|  |  | uninfected-3Δ | 0.147 |  | 0.319 | 0.129 |  | 0.797 |  | 0.052 |  | 0.092 |
|  |  | WT-Δ*invA* | 0.941 |  | 0.492 | 0.958 |  | 0.180 |  | 0.169 |  | 0.384 |
|  |  | WT-3Δ | 0.085 |  | 0.913 | 0.417 |  | 0.336 |  | 0.282 |  | 0.058 |
|  |  | Δ*invA*-3Δ | 0.138 |  | 0.443 | 0.437 |  | 0.177 |  | 0.572 |  | 0.338 |
|  | Infected labeled cells | WT-Δ*invA* | 0.775 |  | 0.608 | 0.681 |  | 0.251 |  | 0.398 |  | 0.380 |
|  |  | WT-3Δ | 0.789 |  | 0.635 | 0.392 |  | 0.477 |  | 0.266 |  | 0.479 |
|  |  | Δ*invA*-3Δ | 0.934 |  | 0.934 | 0.397 |  | 0.554 |  | 0.338 |  | 0.643 |
| Aortic vessels | Labeled cells | uninfected-WT | **0.035** | ***** | 0.300 | 0.457 |  | 0.452 |  | 0.346 |  | 0.677 |
|  |  | uninfected-Δ*invA* | 0.076 |  | 0.192 | 0.269 |  | 0.776 |  | 0.088 |  | 0.606 |
|  |  | uninfected-3Δ | **0.043** | ***** | 0.271 | 0.325 |  | 0.518 |  | 0.213 |  | 0.078 |
|  |  | WT-Δ*invA* | 0.428 |  | 0.406 | 0.201 |  | 0.674 |  | 0.069 |  | 0.915 |
|  |  | WT-3Δ | 0.745 |  | 0.185 | 0.228 |  | 0.285 |  | 0.142 |  | 0.243 |
|  |  | Δ*invA*-3Δ | 0.321 |  | 0.320 | 0.329 |  | 0.440 |  | 0.705 |  | 0.309 |
|  | Infected labeled cells | WT-Δ*invA* | 0.395 |  | 0.364 | 0.303 |  | 0.605 |  | 0.200 |  | 0.447 |
|  |  | WT-3Δ | 0.893 |  | 0.287 | 0.710 |  | 0.905 |  | 0.509 |  | 0.414 |
|  |  | Δ*invA*-3Δ | 0.321 |  | 0.977 | 0.476 |  | 0.632 |  | 0.360 |  | 0.140 |
| Gall bladder | Labeled cells | uninfected-WT | 0.491 |  | 0.226 | 0.727 |  | 0.272 |  | 0.337 |  |  |
|  |  | uninfected-Δ*invA* | 0.392 |  | 0.230 | 0.271 |  | 0.357 |  | 0.177 |  |  |
|  |  | uninfected-3Δ | 0.245 |  | 0.498 | 0.152 |  | 0.578 |  | 0.566 |  |  |
|  |  | WT-Δ*invA* | 0.200 |  | 0.963 | 0.100 |  | 0.704 |  | 0.834 |  |  |
|  |  | WT-3Δ | 0.322 |  | 0.236 | 0.287 |  | 0.650 |  | 0.717 |  |  |
|  |  | Δ*invA*-3Δ | 0.164 |  | 0.268 | **0.041** | ***** | 0.856 |  | 0.481 |  |  |
|  | Infected labeled cells | WT-ΔinvA | **0.020** | ***** | 0.310 | 0.145 |  | 0.642 |  | 0.064 |  |  |
|  |  | WT-3Δ | 0.853 |  | 0.572 | 0.110 |  | 0.409 |  | 0.558 |  |  |
|  |  | ΔinvA-3Δ | 0.134 |  | 0.448 | 0.323 |  | 0.777 |  | 0.091 |  |  |

Asymptotic two-sample Fisher-Pitman permutation tests (One-Way-Test) of the percentages of the labeled cells or of the percentages of the infected labeled cells were performed with the software R, package Rcmdr version 2.5.3 (2019-05-06).

p-values are reported and significance was * p<0.05 (http://www.r-project.org, http://socserv.socsci.mcmaster.ca/jfox/Misc/Rcmdr/).
