## Supplemental Table 2 for "A large panel of chicken cells are invaded *in vivo* by *Salmonella* Typhimurium even when depleted of all known invasion factors"

S2 Table. Antibodies used for cellular characterization of cell subpopulations identified by flow cytometry

| **Target cells** | **Antigen** | **Antibody clone** | **Isotype** | Source | **Secondary antibody** | Source |
| --- | --- | --- | --- | --- | --- | --- |
| Monocytes, macrophages |  | Kul01 | IgG1 | Bio-Rad Cat# MCA5770, RRID:AB_10841619 | Alexa Fluor 488 goat anti-mouse IgG | Invitrogen  (A11017) |
| B Lymphocytes | BU-1 | AV20 | IgG1* | SouthernBiotech Cat# 8395-02, RRID:AB_2796542 |  |  |
| T Lymphocytes | CD3 | CT-3 | IgG1* | SouthernBiotech Cat# 8200-02, RRID:AB_2796418 |  |  |
| Thrombocytes | CD41/CD61  GIIb-IIIa | 11C3 | IgG1 | Bio-Rad Cat# MCA2240, RRID:AB_2128894 | Alexa Fluor 488 goat anti-mouse IgG | Invitrogen  (A11017) |
| Epithelial cells | L-CAM | Anti-L-CAM | IgG1 | DSHB Cat# 7d6, RRID:AB_528115 | Alexa Fluor 488 goat anti-mouse IgG | Invitrogen  (A11017) |
| Endothelial cells | VE-cadherin | Anti-VE-cadherin | IgG | Abcam Cat# ab33168, RRID:AB_870662 | Alexa Fluor 488 donkey anti-rabbit IgG | Invitrogen  (A21206) |

*Conjugated FITC

| **Target cells** | **Control** | Source |
| --- | --- | --- |
| Monocytes, macrophages | Mouse IgG1-Alexa Fluor 488 | SouthernBiotech Cat# 0102-30, RRID:AB_2793862 |
| B Lymphocytes | Mouse IgG1-FITC | SouthernBiotech Cat# 0102-02, RRID:AB_2793846 |
| T Lymphocytes | Mouse IgG1-FITC | SouthernBiotech Cat# 0102-02, RRID:AB_2793846 |
| Thrombocytes | Mouse IgG1-Alexa Fluor 488 | SouthernBiotech Cat# 0102-30, RRID:AB_2793862 |
| Epithelial cells | Mouse IgG1-Alexa Fluor 488 | SouthernBiotech Cat# 0102-30, RRID:AB_2793862 |
| Endothelial cells | Normal rabbit IgG | Santa Cruz Biotechnology Cat# sc-3888, RRID:AB_737196 |
